# p27 regulates the autophagy-lysosomal pathway via the control of Ragulator and mTOR activity in amino acid deprived cells

**DOI:** 10.1101/2020.01.07.896860

**Authors:** Ada Nowosad, Pauline Jeannot, Caroline Callot, Justine Creff, Renaud T. Perchey, Carine Joffre, Patrice Codogno, Stephane Manenti, Arnaud Besson

**Author notes:** Corresponding author: Arnaud Besson, LBCMCP UMR5088 CNRS - Université Paul Sabatier, Batiment 4R3b1, 118 route de Narbonne, 31062 Toulouse cedex 9, France.

## Abstract

Autophagy is a catabolic process whereby cytoplasmic components are degraded within lysosomes, allowing cells to maintain energy homeostasis during nutrient depletion. Several studies have shown that the CDK inhibitor p27^Kip1^ promotes starvation-induced autophagy. However, the underlying mechanism remains unknown. Here, we report that in amino acid deprived cells, p27 controls autophagy via an mTORC1-dependent mechanism. During prolonged amino acid starvation, a fraction of p27 is recruited to lysosomes where it interacts with LAMTOR1, a component of the Ragulator complex required for mTORC1 lysosomal localization and activation. p27 binding to LAMTOR1 prevents Ragulator assembly and function and subsequent mTORC1 activation, thereby promoting autophagy. Conversely, upon amino acid withdrawal, p27^−/−^ cells exhibit elevated mTORC1 signaling, impaired lysosomal activity and autophagy, and resistance to apoptosis. This is associated with sequestration of TFEB in the cytoplasm, preventing the induction of lysosomal genes required for lysosomal function. Silencing of LAMTOR1 or mTOR inhibition restores autophagy and induces apoptosis in p27^−/−^ cells. Together, these results reveal a direct, coordinated regulation between the cell cycle and cell growth machineries.

## INTRODUCTION

In all organisms, cell growth is coupled to cell division to allow normal development and maintain homeostasis. How cells coordinate the machinery that regulates cell growth, at the heart of which lies the mTOR kinase, with the machinery that controls cell division, driven by cyclin/CDK complexes, has been the subject of considerable interest for several decades^1^. This coordination has been mostly studied under normal metabolic conditions and except for notable exceptions, such as early embryonic development, it appears that growth control drives the activity of the cell cycle machinery. Here we investigated this question in conditions of metabolic restriction to determine if the cell cycle machinery could in turn regulate the growth control machinery.

p27^Kip1^ (p27) was initially identified as a cyclin/CDK inhibitor^2, 3^. Due to its ability to induce cell cycle arrest, p27 acts as a tumor suppressor and p27^−/−^ mice display multiple organ hyperplasia and spontaneously develop pituitary tumors^4^. Nevertheless, in contrast to classic tumor suppressors such as p53 or Rb, inactivating mutations of the p27 gene are extremely rare and its inactivation in cancer is rather caused by enhanced degradation, attenuated transcription or translation, or mislocalization in the cytoplasm^2, 3, 5^. The latter correlates with poor prognosis in a variety of cancers, suggesting a direct contribution of cytoplasmic p27 to tumor progression^2, 3^. In fact, knock-in mice in which p27 is largely sequestered in the nucleus due to defective export to the cytoplasm (p27^S10A^) are partially resistant to tumorigenesis^6^. Conversely, another knock-in model in which p27 cannot bind to cyclin-CDKs (p27^CK−^) revealed that p27 can act as an oncogene, as these mice display an increased susceptibility to both spontaneous and induced tumorigenesis compared to p27^−/−^ mice^7, 8^. How exactly p27 acts as an oncogene remains elusive, but could be due to the regulation of several other cellular processes by p27. Indeed, p27 has been involved in the control, sometimes in a CDK-independent manner, of cell migration and invasion, differentiation, cytokinesis, transcriptional repression, apoptosis and autophagy^2, 9–15^.

Autophagy is a catabolic process by which intracellular components are degraded and recycled by the lysosomal machinery^16, 17^. Depending on the pathway implicated in the delivery of cargo for degradation, there are three major types of autophagy: macroautophagy (hereafter referred to as autophagy), microautophagy and chaperone-mediated autophagy^17^. Autophagic degradation begins with formation of double-membrane autophagosomes that engulf the cytoplasmic material destined for elimination. Autophagosomes then fuse with the endocytic compartment and eventually deliver their cargo to lysosomes for degradation by lysosomal enzymes^17, 18^. Following proteolysis, the resulting molecules are released back to the cytosol through lysosomal permeases for reuse^19^. Although autophagy occurs constantly in cells and is required to control the quality of cytoplasm, its levels dramatically increase in stress situations, such as nutrient withdrawal, and particularly under amino acid (aa) deprivation^20^. In these conditions, autophagy recycles cellular components and the products of proteolysis are delivered back to the cytoplasm, where they are used as energy source or as building blocks for protein synthesis^17^.

Autophagy is negatively modulated by mTORC1, a master regulator of cell growth^21, 22^. mTORC1 is a multi-protein complex consisting of mTOR and its regulatory binding partners Raptor, MLST8, PRAS40 and Deptor^22^. The main function of mTORC1 is to orchestrate the balance between anabolic and catabolic metabolism. Via a complex network of nutrient sensors, in presence of aa, mTORC1 is recruited to lysosomal membranes by Rag GTPases, themselves anchored by the Ragulator complex, where the kinase activity of mTORC1 can be stimulated by the GTPase Rheb that also resides on lysosomes^23–26^. When nutrients are abundant, mTORC1 promotes protein synthesis via phosphorylation of substrates implicated in the regulation of translation, such as S6 kinases and 4E-binding proteins. At the same time, mTORC1 inhibits autophagy at multiple levels by phosphorylating and inactivating proteins involved in autophagosome formation (such as ULK1, AMBRA1, Atg13 and Atg14) and maturation (UVRAG)^21, 22, 27, 28^. mTORC1 also phosphorylates TFEB, a transcription factor that drives the expression of many pro-autophagic and lysosomal genes, on S142 and S211, resulting in TFEB cytoplasmic retention, thereby inhibiting its activity^29–31^. Conversely, under nutrient shortage, mTOR is inactivated, allowing the autophagic machinery to form autophagosomes. During prolonged aa starvation, partial mTORC1 reactivation triggers the reformation of lysosomes from autolysosomes via a process called autophagic lysosome reformation (ALR)^32^. Thus, the role of mTORC1 in autophagy is complex and context dependent: while mTORC1 inhibition is required for autophagy initiation, its cyclic reactivation by autophagy-generated nutrients allows maintaining autophagy during prolonged starvation by restoring the lysosomal population.

Cytoplasmic p27 was recently described as a positive regulator of basal and starvation-induced autophagy and to protect cells in conditions of metabolic stress from apoptosis by promoting autophagy^14, 33–37^. Glucose or serum deprivation activates the energy/nutrient sensing kinases LKB1 and AMPK, in turn AMPK phosphorylates p27 on S83, T170 and T198, causing its stabilization and cytoplasmic retention^14, 36, 38^. Loss of Tsc2, a subunit of Tuberous Sclerosis Complex that acts as a GTPase activating protein (GAP) for Rheb, also causes AMPK activation and promotes p27-dependent autophagy^36–39^. Expression of a p27 T198D mutant that localizes in the cytoplasm is sufficient to increase basal autophagy, while p27 silencing interferes with autophagy induction upon serum or glucose deprivation and causes apoptosis^14^. However, while upstream events causing p27 cytoplasmic localization in response to metabolic stress have been studied in detail, the molecular mechanism underlying the pro-autophagic role of p27 remains completely unknown^35^.

Herein, we investigated the mechanism by which p27 modulates autophagy upon aa starvation and found that a fraction of p27 relocalizes to the lysosomal compartment where it binds to the Ragulator subunit LAMTOR1 and participates in the inhibition of mTOR, allowing maintenance of autophagy. In absence of p27, increased mTORC1 activity results in TFEB cytoplasmic retention, decreased lysosomal function and reduced autophagic flux in response to aa deprivation and promotes cell survival. These results indicate that upon prolonged aa withdrawal, the cell cycle inhibitor p27 exerts a direct negative feedback on the master cellular growth regulator mTOR by participating in its inhibition, illustrating the crosstalk between the cell division and cell growth machineries.

## RESULTS

### p27 promotes autophagy flux in amino acid-deprived cells

p27 was shown to promote autophagy in glucose-starved cells^14^. While the impact of glucose deprivation on autophagy is still a matter of debate^40, 41^, aa starvation is the most potent and best characterized autophagy inducer^20, 42^. To determine the effect of p27 on autophagy upon aa withdrawal, levels of the autophagosome marker LC3B-II^43, 44^ were measured in p27^+/+^ and p27^−/−^ HPV E6-immortalized mouse embryo fibroblasts (MEFs) by immunoblot (Fig. 1A, B). Whereas LC3B-II levels initially decreased similarly in p27^+/+^ and p27^−/−^ MEFs, suggesting that autophagy induction occurred normally in absence of p27, they were consistently elevated in p27^−/−^ cells compared to p27^+/+^ cells during prolonged aa starvation (Fig. 1A, B). This difference was statistically significant at 48 h of aa deprivation and this duration was used in most subsequent experiments. LC3B immunostaining confirmed these observations, with p27^−/−^ cells exhibiting an increased number of LC3B puncta after 48 h of starvation (Fig. 1C, D). ULK1 dephosphorylation on S757, targeted by mTORC1^45^, occurred in a similar manner in p27^+/+^ and p27^−/−^ cells and ULK1 expression was progressively downregulated (Fig. S1A), as previously reported^46, 47^. AMPK was not activated by LKB1 on T172 upon aa starvation (Fig. S1A), in agreement with previous studies^48^. Consistent with the lack of AMPK stimulation, p27 phosphorylation on T198 did not increase after aa withdrawal (Fig. S1B), unlike what was observed previously under glucose and/or serum starvation^14^.

**Figure 1:**
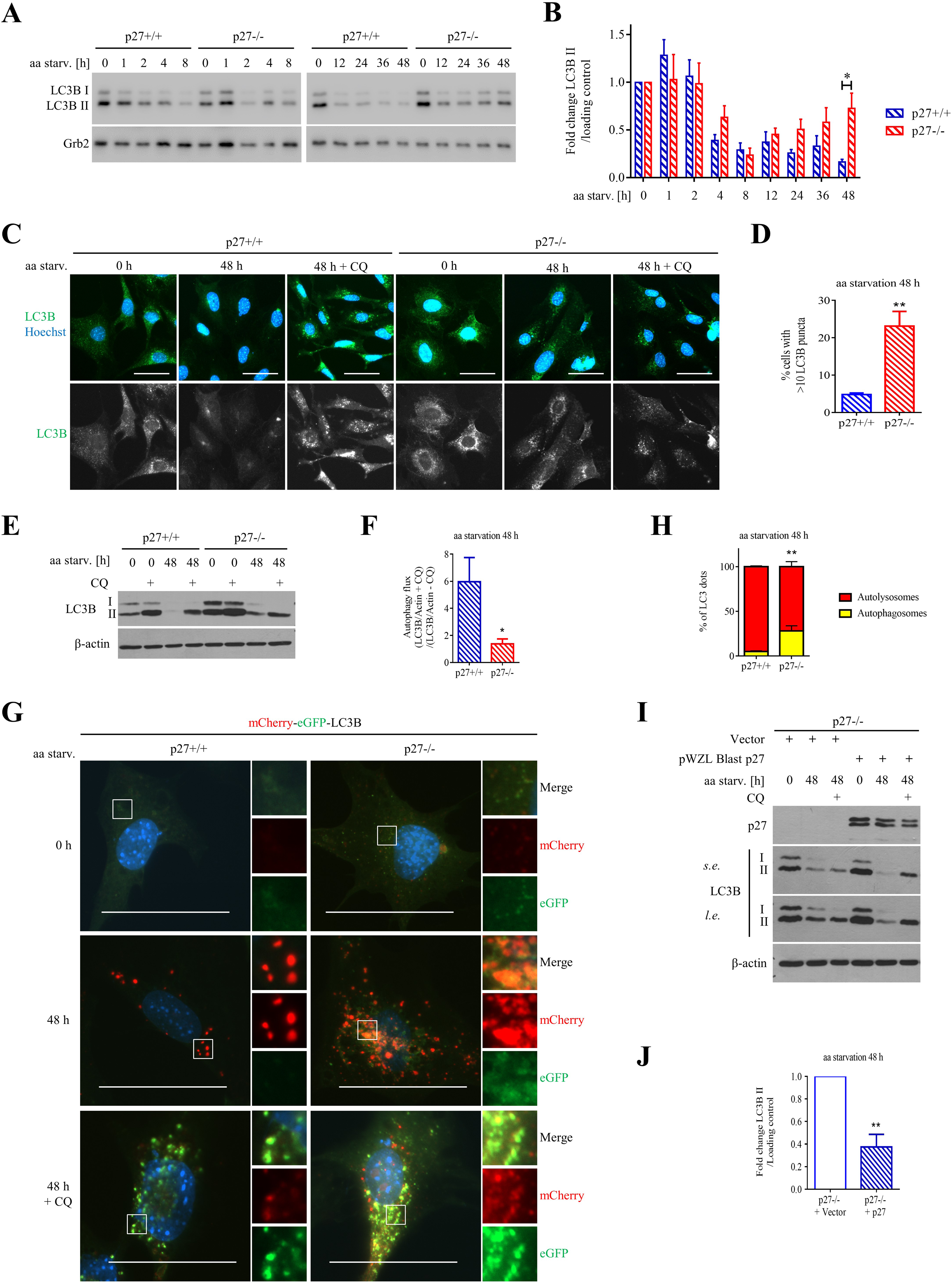
p27 promotes autophagy flux during prolonged amino acid starvation. **(A)** LC3B immunoblotting in p27^+/+^ and p27^−/−^ MEFs in full medium (0 h) or aa-deprived for the indicated times. Grb2 was used as loading control. **(B)** Densitometry analysis from at least 3 experiments of the ratio of LC3B-II levels by that of loading control in p27^+/+^ and p27^−/−^ MEFs as in A, normalized to 0 h. **(C)** LC3B immunostaining in p27^+/+^ and p27^−/−^ MEFs in full medium (0 h) or aa-deprived for 48 h ± 50 µM CQ for 2 h. Scale bars are 50 µm. **(D)** Percentage of p27^+/+^ and p27^−/−^ MEFs with >10 LC3B puncta after 48 h of aa starvation. At least 100 cells were analyzed per genotype in each experiment (n=5). **(E)** LC3B immunoblotting in p27^+/+^ and p27^−/−^ MEFs in full medium (0 h) or aa-deprived for 48 h ± 50 µM CQ for the last 2 h. β-actin was used as loading control. **(F)** LC3B-II turnover, which corresponds to the ratio of (LC3B-II+CQ/β-actin)/(LC3B-II-CQ/β-actin), as in E, from 11 experiments. **(G)** Images of LC3B puncta in p27^+/+^ and p27^−/−^ cells expressing mCherry-eGFP-LC3B in full medium or aa-starved for 48 h ± 50 µM CQ for 2 h. Autophagosomes appear in yellow and autolysosomes in red. Scale bars are 50 µm. Levels of mCherry-eGFP-LC3B expression by immunoblot are shown in Fig. S1G. **(H)** Quantification of autophagosomes (yellow) and autolysosomes (red) from cells described in G after 48 h aa starvation from 3 experiments. At least 399 LC3B puncta were analyzed per genotype in each experiment. **(I)** LC3B and p27 immunoblots of p27^−/−^ cells infected with empty or p27 expression vector in full medium (0 h) or aa deprived for 48 h ± 50 µM CQ for 2 h. ‘s.e.’ = short exposure; ‘l.e.’ = long exposure. β-actin was used as loading control. **(J)** Densitometry analysis of the ratio of LC3B-II levels by that of loading control after 48 h aa deprivation in cells described in I, normalized to p27^−/−^ + empty vector cells, from 4 experiments. **(B, D, F, H, J)** Bar graphs show means ± SEM. Statistical significance was evaluated by multiple t test (B), unpaired t-test with Welch’s corrections (D, J), or by 2-way ANOVA test (F, H); **: p ≤ 0.01; *: p ≤ 0.05.

LC3B-II levels transiently increase following phagophore formation and then decrease during autophagosome maturation, as LC3B-II located on autophagosome inner and outer membranes is degraded or cleaved, respectively^43, 44^. To distinguish between autophagy induction and the block of late stage autophagy, which also results in LC3B-II accumulation, autophagy flux was evaluated by several methods. First, LC3B degradation in autolysosomes was inhibited with the lysosomotropic alkalizing agent Chloroquine (CQ). CQ treatment revealed that while autophagy flux was similar in p27^+/+^ and p27^−/−^ MEFs in short-term starvation (Fig. S1C, D), it was markedly reduced in p27^−/−^ MEFs compared to p27^+/+^ cells during prolonged aa starvation (Fig. 1E, F). Second, p62/SQSTM1, which accumulates in cells with impaired autophagy flux^43, 44, 49^, was elevated in aa starved p27^−/−^ cells compared to wild-type (Fig. S1E, F). Third, autophagosome maturation was monitored using p27^+/+^ and p27^−/−^ MEFs stably expressing tandem mCherry-eGFP-LC3B (Fig. S1G) that labels autophagosomes in yellow and autolysosomes in red due to quenching of eGFP fluorescence in the acidic lysosomal environment^43, 44, 50^. Under basal conditions, LC3B signal was diffuse in the cytoplasm and aa starvation induced the formation of fluorescent LC3B puncta, however, p27^−/−^ cells had a decreased fraction of autolysosomes (red) compared to p27^+/+^ cells (72% vs 95%) (Fig. 1G, H), indicating that p27 promotes autophagosome maturation. Finally, re-expression of p27 in p27^−/−^ cells by retroviral infection restored LC3B degradation upon aa starvation (Fig. 1I, J), confirming the involvement of p27 in this process.

The role of p27 in promoting autophagy has been associated with localization of p27 in the cytoplasm and with its capacity to inhibit CDK activity^14, 36^. To test whether these features were also needed in aa deprivation conditions, we measured autophagy flux in p27^S10A^ MEFs, in which p27 is sequestered in the nucleus due to impaired nuclear export, and in p27^CK−^ MEFs, in which p27 cannot bind to and inhibit cyclin-CDK complexes^6, 7^. On aa deprivation, p27^S10A^ cells behaved like p27^−/−^ cells and had decreased autophagy flux. In contrast, p27^CK−^ had the same pro-autophagic properties as wild-type p27 (Fig. S1H, I). In line with these results, p27^CK−^ expression in p27^−/−^ cells restored autophagy flux (Fig. S1J, K), as observed for wild-type p27 (Fig. 1I, J). Taken together, our data indicate that p27 promotes autophagy flux in aa-deprived cells and this role requires p27 cytoplasmic localization but is independent of cyclin/CDK inhibition.

### A fraction of p27 localizes to lysosomal compartments during amino acid deprivation

Previous data showed that autophagy induction by glucose deprivation promotes p27 cytoplasmic localization^14^. Proximity ligation assays (PLA) using antibodies against the lysosome marker LAMP1 and p27 indicated that a small fraction of p27 localized to the lysosomal compartment and this was increased following aa starvation (Fig. 2A, B and S2). However, p27 did not interact directly with LAMP1, as tested by pull-downs using recombinant proteins (data not shown). These results were confirmed using a subcellular fractionation approach to enrich the lysosomal compartment^51^, evidenced by the presence of LAMP2 and LAMTOR1, two proteins that localize to lysosomal membranes (Fig. 2C). A small amount of p27 was detected in the lysosomal fraction, which increased following aa starvation for 18 h. Conversely, the amount of active p70 S6K1 and mTOR in the lysosomal fraction decreased following aa starvation (Fig. 2C), as expected^23^. These results suggest that a fraction of p27 localizes to lysosomes upon aa withdrawal, where it may regulate autophagy.

**Figure 2:**
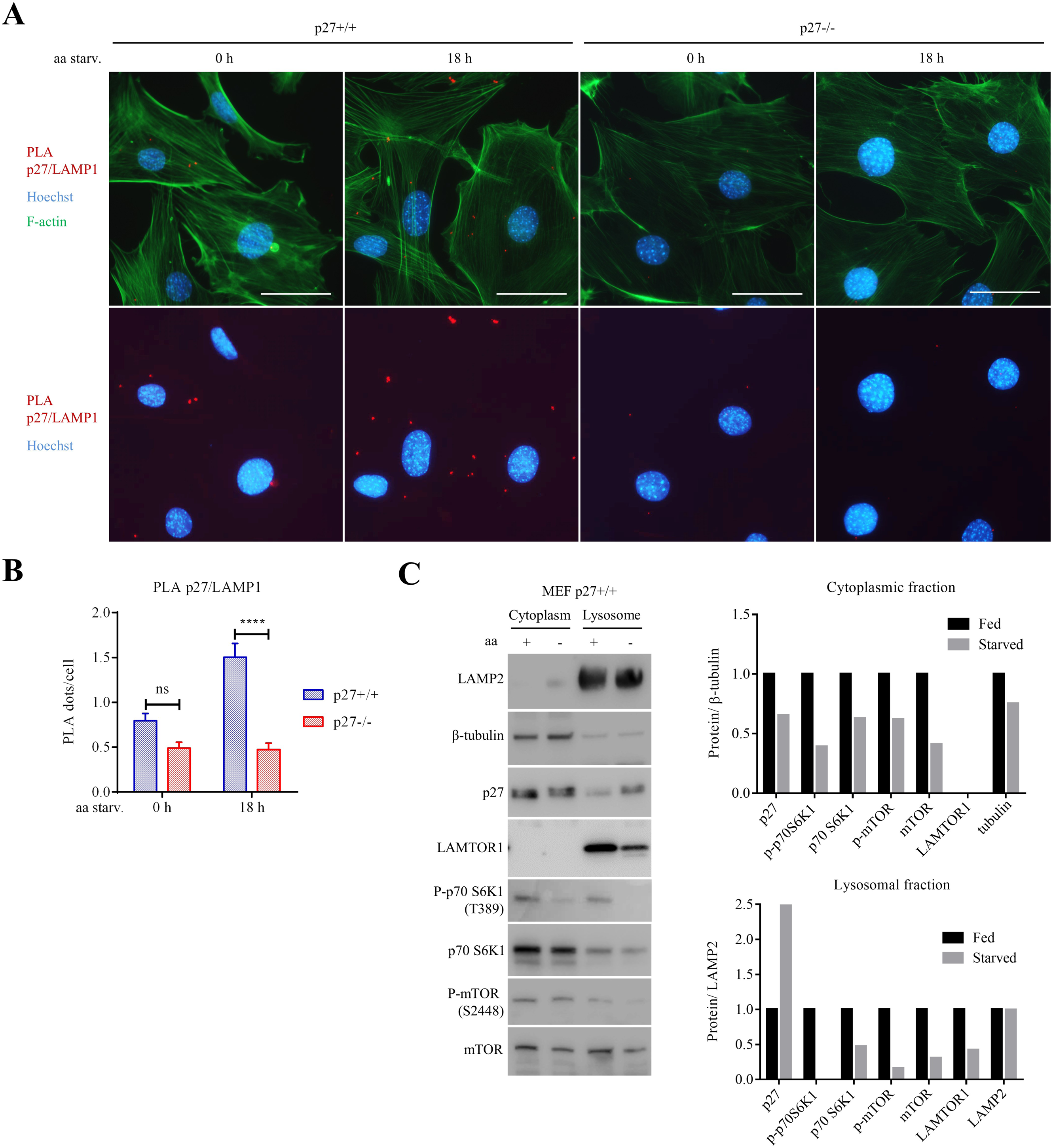
p27 localizes on lysosomes upon amino acid starvation. **(A)** Proximity ligation assays (PLA) for p27 and the lysosomal marker LAMP1 in p27^+/+^ MEFs in full medium or aa deprived for 18 h. p27^−/−^ MEFs were used as negative control. F-actin was stained with phalloïdin. Scale bar are 50 µm. Controls PLA reactions with single antibody or without primary antibodies are shown in Fig. S2. **(B)** Bar graph shows the mean number of PLA dots per cell ± SEM as described in A (n cells counted: p27^+/+^ 0 h=1618, p27^+/+^ 18 h=1797, p27^−/−^ 0 h=899, p27^−/−^ 18 h=239). Statistical significance was evaluated by 2-way ANOVA test; ****: p ≤ 0.0001. **(C)** Immunoblots for the indicated proteins in cytoplasmic and lysosome-enriched fractions prepared from p27^+/+^ MEFs in full medium or aa-deprived for 18 h. Graphs show the ratio of densitometry analysis for each protein to β-tubulin (cytoplasmic fractions, top graph) or to LAMP2 (lysosomal fractions, bottom graph) normalized to the corresponding full medium condition.

### p27 promotes autophagosome maturation

The fusion of autophagosomes with lysosomes plays an essential role in autophagic cargo degradation^52^. To further dissect the role of p27 in autophagy, the morphology of LC3B vesicles, which include pre-autophagosomal structures, autophagosomes and autolysosomes, was examined in p27^+/+^ and p27^−/−^ aa-starved MEFs. During prolonged aa starvation, LC3B-positive vesicles formed large ring-shaped aggregates mainly in the perinuclear zone of p27^+/+^ cells (Fig. S3A, B), previously identified as intermediate structures in the process of autophagosome maturation^53^. These were rarely observed in p27^−/−^ cells, in which small LC3-positive vesicles were abundant instead (Fig. S3A, B).

The autophagy receptor p62/SQSTM1 is recruited to autophagosomes prior to their closing and degraded along with its cargo in autolysosomes^54^. Interestingly, ring-like structures in p27^+/+^ MEFs rarely colocalized with p62, while small LC3-positive vesicles in p27^−/−^ cells exhibited frequent p62 colocalization (Fig. S3A, C). These observations indicate that recognition and sequestration of autophagosome cargo is not affected in cells lacking p27 since p62, which targets cargo to LC3B+ autophagosomes, colocalizes with LC3B^54, 55^. However, the persistence of p62/LC3B colocalization in p27^−/−^ cells suggests defective p62 degradation, which could be due to failure of autophagosome/lysosome fusion or reduced proteolytic activity within autolysosomes. Indeed, the loss of p62 within ring-shaped structures in p27^+/+^ MEFs supports the idea that they represent mature autolysosomes with partially degraded cytoplasmic material, including p62. This was confirmed by CQ treatment, which blocks autophagic degradation but not autophagosome maturation. CQ restored p62 and LC3B colocalization in p27^+/+^ cells, without noticeable effect in p27^−/−^ cells (Fig. S3A, C).

Autophagosome/lysosome fusion was not affected in p27^−/−^ cells since colocalization of LC3B (autophagosome marker) and LAMP2 (lysosomal marker) was similar in aa-deprived p27^+/+^ and p27^−/−^ cells treated with CQ to prevent LC3B degradation (Fig. S3D, E). Thus, our results suggest that p27 controls autophagy after autophagosome/lysosome fusion, possibly by regulating lysosome function.

### Lysosomal function is decreased in absence of p27

To investigate the role of p27 in lysosomal function, the degradative capacity of lysosomes in p27^+/+^ and p27^−/−^ MEFs was compared (Fig. 3). First, BSA dequenching assays were performed, in which self-quenched BODIPY-FITC BSA (DQ-BSA) acts as a lysosomal proteolysis sensor when BODIPY fluorescence is dequenched by protease activity within lysosomes^56^. These experiments showed that BODIPY signal was abundant in p27^+/+^ cells in aa-deprivation conditions as a result of BSA dequenching, whereas only low signal was detected in p27^−/−^ cells (Fig. 3A, B). CQ, which delays BSA dequenching by suppressing lysosomal function, was used as a negative control in these experiments (Fig. 3A). This difference was not due to altered endocytosis in p27^−/−^cells, since uptake of TRITC-Dextran 40, which also enters cells via fluid-phase endocytosis^57^, was similar in p27^+/+^ and p27^−/−^ cells (Fig. S3F).

**Figure 3:**
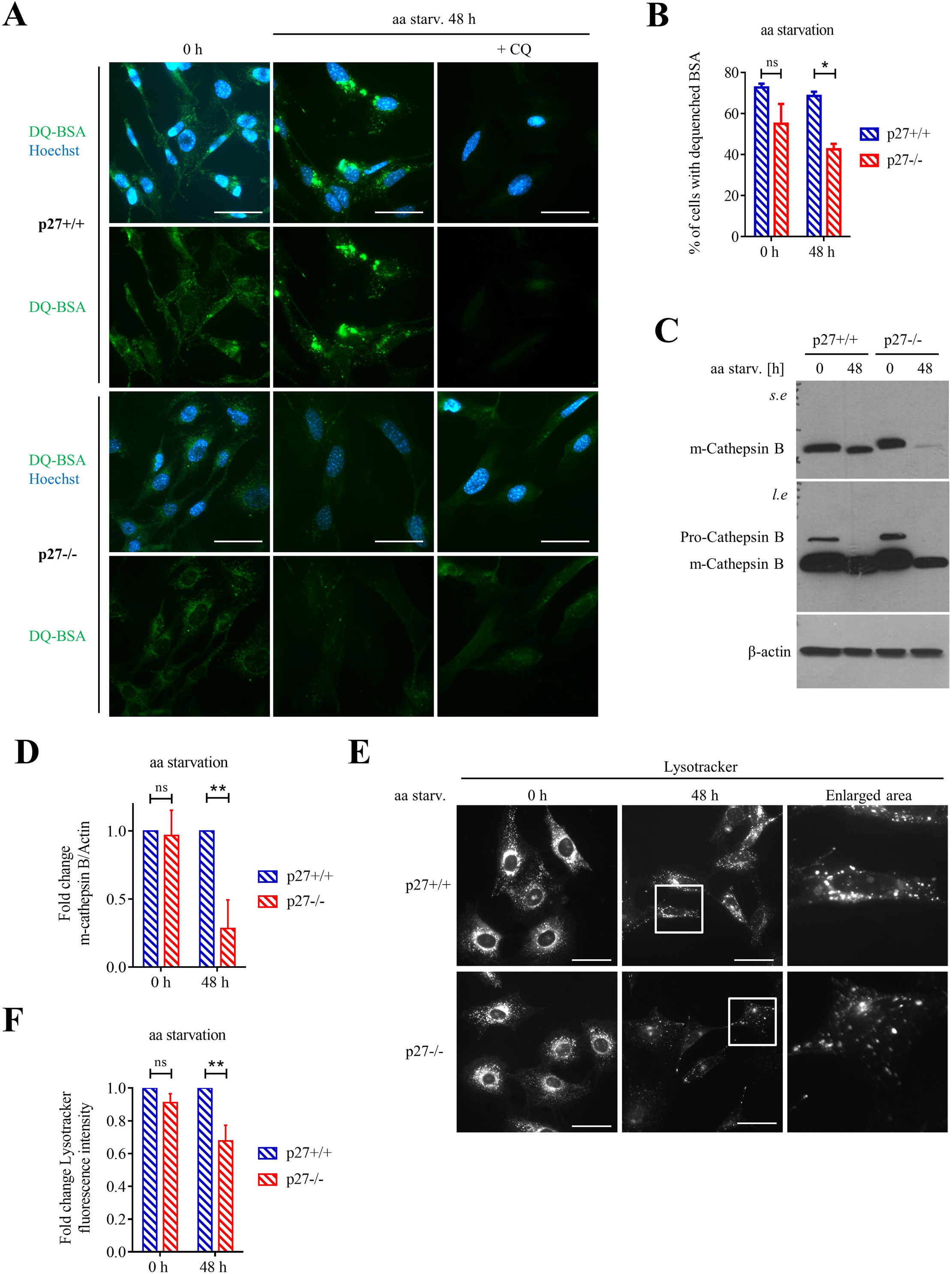
p27 loss decreases lysosomal function. **(A)** Images of DQ-BSA in p27^+/+^ and p27^−/−^ MEFs in full medium or aa-starved for 48 h. CQ 50 µM for 2 h was used as negative control. Scale bar are 50 µm. **(B)** Quantification of the percentage of p27^+/+^ and p27^−/−^ MEFs exhibiting dequenched BSA signal in full medium and after 48 h aa starvation from 3 experiments. At least 81 cells per condition per experiment were analyzed. **(C)** Cathepsin-B immunoblot in cells in full medium or aa-starved for 48 h. Upper bands represent pro-Cathepsin B and the lower bands correspond to mature m-Cathepsin B. ‘s.e.’ = short exposure; ‘l.e.’ = long exposure. β-actin was used as loading control. **(D)** Densitometry analysis of m-cathepsin B/actin ratio from immunoblots described in C normalized to p27^+/+^ cells in each condition from 3 experiments. **(E)** Lysotracker (100 nM) staining in p27^+/+^ and p27^−/−^ MEFs in full medium or aa-starved for 48 h. Scale bar are 50 µm. **(F)** Quantification of Lysotracker signal intensity in p27^+/+^ and p27^−/−^ MEFs as described in E normalized to p27^+/+^ cells in each condition from 3 experiments. At least 10 images of each genotype, acquired with a 20X objective, were analyzed per condition per experiment. **(B, D, F)** Bar graphs show means ± SEM. Statistical significance was evaluated by 2-way ANOVA; ** p ≤ 0.01; *: p ≤ 0.05.

Second, Cathepsin B, a lysosomal enzyme essential for autophagy, is synthesized as inactive zymogen (pro-cathepsin B, 44 kDa) and is activated by proteolytic cleavage into its mature forms (m-cathepsin B, 24 and 27 kDa) in the acidic lysosomal environment^58^. m-cathepsin B levels were decreased in aa-starved p27^−/−^ MEFs compared to p27^+/+^ cells (Fig. 3C, D), suggesting that proteolytic activity in lysosomes is reduced in absence of p27. Surprisingly, there was no compensatory elevation of pro-cathepsin B levels in p27^−/−^ cells, implying that expression of the protein is affected in aa-deprivation conditions in these cells (see below).

Finally, while Lysotracker staining of acidic vesicles in p27^+/+^ and p27^−/−^ cells was similar in full medium, it was decreased in aa-starved p27^−/−^ cells compared to wild-type (Fig. 3E, F). This decrease was not due to a reduction of lysosomal compartment size, estimated by LAMP2 immunostaining (Fig. S3G, H). These results indicate that p27 regulates lysosomal acidification and the activation of lysosomal proteases, thereby affecting autophagic degradation.

### p27 affects Ragulator assembly and function on lysosomal membranes

Our results suggest that p27 controls autophagy by acting directly from the lysosomal surface. Interestingly, p27RF-Rho (p27^Kip1^ Releasing Factor from RhoA), a previously identified p27 binding partner^59^, is in fact p18/LAMTOR1 (Late ensodome/lysosome membrane adaptor, MAPK and mTORC1). LAMTOR1 has been implicated in trafficking of intracellular organelles, lysosomal function, cytoskeleton regulation and modulation of mTOR and MAPK signaling^23, 59–62^. LAMTOR1 acts as a lysosomal anchor and scaffold for the other subunits of the pentameric Ragulator complex, which mediates mTORC1 activation^23, 63–65^. In response to aa, Ragulator acts as a platform to recruit Rag GTPases to lysosomal. In turn Rags recruit mTOR on lysosomes, where it can be activated by Rheb^23, 24, 62, 66–68^. Ragulator and SLC38A9 also act as atypical GEFs for RagC/D and A/B, respectively, and GTP loading of RagA/B is required for mTOR recruitment to lysosomes^69^. Thus, LAMTOR1 plays an essential role in the assembly and localization of Ragulator and in the recruitment of mTOR on lysosomes and LAMTOR1 depletion impairs lysosome maturation, fusion with autophagosomes and autophagy flux^60, 61, 70^. Here, we investigated the role of the p27/LAMTOR1 interaction in the context of autophagy.

The p27/LAMTOR1 interaction was confirmed by co-immunoprecipitation (co-IP) in HEK293 cells overexpressing LAMTOR1 and p27 or p27^CK−^ (Fig. 4A), indicating that this interaction does not require cyclin-CDK complexes. Pull-down assays with recombinant LAMTOR1 and p27 or p27^CK−^ showed a direct interaction between the two partners (Fig. 4B). Pull-downs with various GST-p27 mutants on HEK293 lysates expressing LAMTOR1 indicated that LAMTOR1 bound to the C-terminal half of p27 (aa 88-198), while binding to a p27^1–190^ mutant was decreased (Fig. 4C). This is consistent with the C-terminus of p27 being required for binding to RhoA and that LAMTOR1 competes with RhoA for binding to p27^13, 59, 71^. The interaction of LAMTOR1 with p27 was confirmed on endogenous proteins by co-IP in U251N cells (Fig. 4D) as well as by PLA in MEFs (Fig. 4E, F and S2). Importantly, while only few PLA dots were visible in full medium, the p27/LAMTOR1 interaction sharply increased in aa-starved p27^+/+^ cells (Fig. 4E, F).

**Figure 4:**
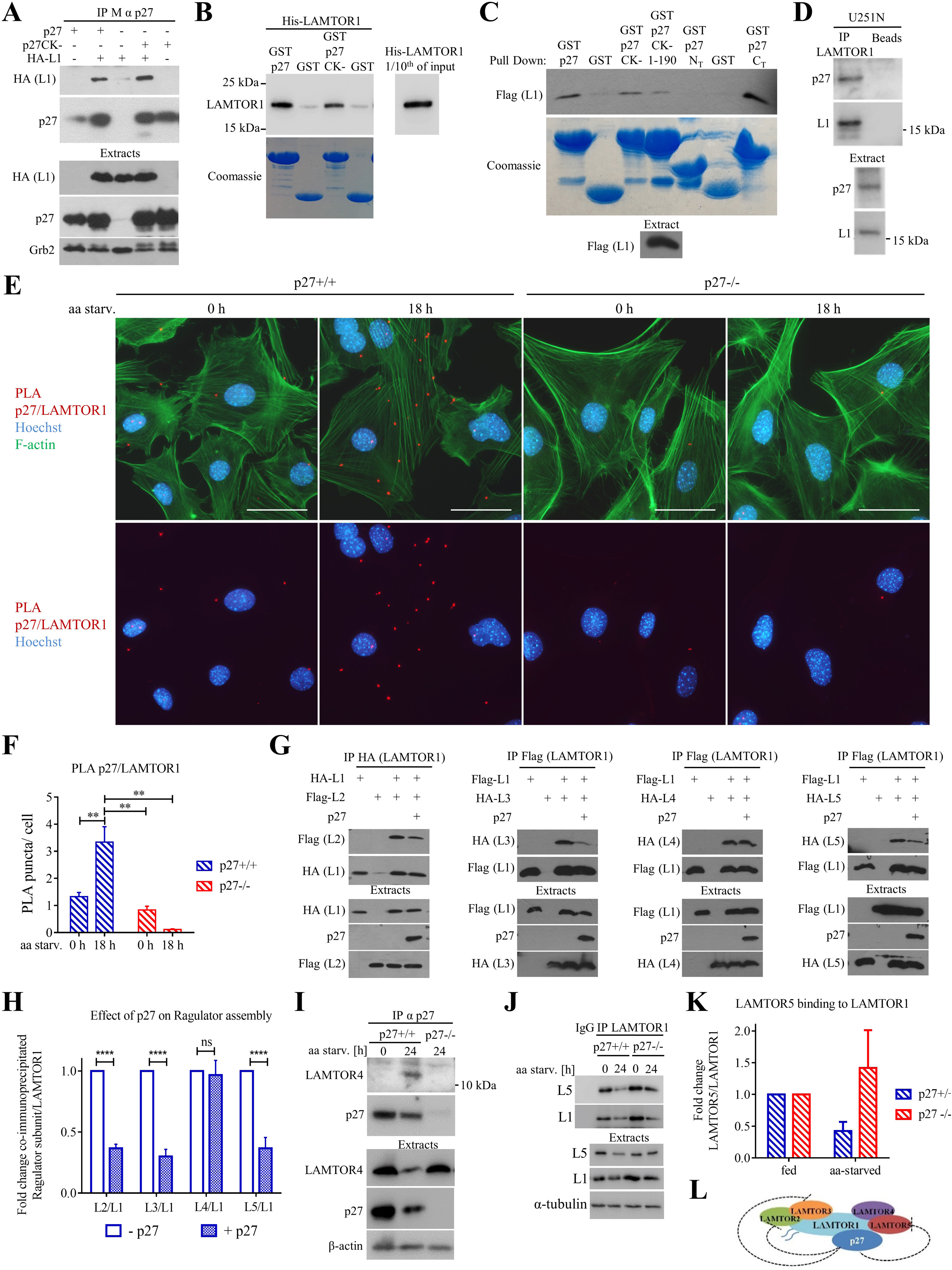
p27 binds to LAMTOR1 and inhibits Ragulator assembly. **(A)** p27 was immunoprecipitated from HEK293 cells transfected with p27 or p27^CK−^ and/or HA-LAMTOR1 (L1), membranes were probed with anti-HA (LAMTOR1) and p27. Transfected protein levels were determined in extracts with the same antibodies. Grb2 was used as loading control. **(B)** Pull-downs of recombinant His-tagged LAMTOR1 with GST-p27 or GST-p27^CK^-beads, GST only beads were used as control. The amounts of beads used were determined by Coomassie staining. A tenth of His-LAMTOR1 input at the same exposure time is shown on the right. **(C)** HEK293 expressing Flag-LAMTOR1 were subjected to pull-downs using GST beads of various p27 deletion mutants and immunoblotted with anti-Flag (LAMTOR1) antibodies. p27 N_T_ = aa 1-87 and p27 C_T_ = aa 88-198. The amounts of beads used were determined by Coomassie staining. Ectopic expression of LAMTOR1 in cell extract was confirmed by anti-Flag immunoblot. **(D)** Endogenous LAMTOR1 was immunoprecipitated from U251N cell lysates with anti-LAMTOR1 antibodies and blotted against p27 and LAMTOR1. Protein A beads were used as negative control. Levels of endogenous proteins were determined with anti-p27 and anti-LAMTOR1 antibody. **(E)** PLA using p27 and LAMTOR1 antibodies on p27^+/+^ MEFs in full medium or aa deprived for 18 h. p27^−/−^ MEFs were used as negative control. F-actin was stained with phalloïdin. Scale bar are 50 µm. Controls PLA reactions with single antibody or without primary antibodies are shown in Fig. S2. **(F)** Bar graph shows the mean number of PLA dots per cell ± SEM as described in E (n cells counted: p27^+/+^ 0 h=2087, p27^+/+^ 18 h=1677, p27^−/−^ 0 h=579, p27^−/−^ 18 h=225). Statistical significance was evaluated by 2-way ANOVA; **: p ≤ 0.01. **(G)** HEK293 cells were transfected with tagged LAMTOR1 and/or another Ragulator subunit and/or p27. LAMTOR1 co-IPs were probed for the co-transfected Ragulator subunit to determine the impact of p27 expression on the ability of LAMTOR1 to interact with its partners. **(H)** Densitometry analysis of experiments described in G. Signal intensity values of LAMTOR2, −3, − 4 and −5 bound to LAMTOR1 were normalized to that in absence of p27. Bar graph shows means ± SEM from 3 independent experiments. Statistical significance was evaluated by 2-way ANOVA; ns: p> 0.05; ****: p ≤ 0.0001. **(I)** p27 was immunoprecipitated from p27^+/+^ MEFs in full medium or aa starved for 24 h, p27^−/−^ cells aa starved for 24 h were used as negative control. The amount of LAMTOR4 co-precipitated with p27 was determined. Levels of LAMTOR4 and p27 in the corresponding extracts are shown. β-actin was used as loading control. **(J)** LAMTOR1 was immunoprecipitated from p27^+/+^ and p27^−/−^ MEFs in full medium or aa starved for 24 h and the amount of LAMTOR5 co-precipitated was determined. Control immunoprecipitation with rabbit IgG was used as control. Levels of LAMTOR1 and −5 in the corresponding extracts are shown. β-actin and β-tubulin were used as loading control. **(K)** Ratio of LAMTOR5 co-precipitated by that of LAMTOR1 immunoprecipitated from p27^+/+^ and p27^−/−^ MEFs in full medium or aa starved for 24 h, normalized to full medium condition from 2 experiments as described in J. **(L)** Schematic summarizing the impact of p27 on Ragulator assembly.

Since LAMTOR1 acts as a scaffold for the other Ragulator subunits (LAMTOR2, −3, −4 and − 5)^23, 72–74^, we tested if p27 expression affects the interaction of LAMTOR1 with its partners by co-IP in HEK293 cells expressing LAMTOR1 and one of its partners. p27 competed with LAMTOR2, −3 and −5, but not LAMTOR4, for binding to LAMTOR1 (Fig. 4G-H), suggesting that p27 interferes with Ragulator complex assembly. Since LAMTOR4 remains bound to LAMTOR1 in presence of p27, we tried to co-IP endogenous LAMTOR4 and p27 in MEFs. In full medium, no LAMTOR4/p27 interaction was detected, but it became apparent after aa starvation for 24 h (Fig. 4I). No LAMTOR2, −3 and −5 could be detected in these experiments. Similarly, a LAMTOR4/p27 PLA was readily detectable, at low levels in full medium and sharply increased in aa starved MEFs (Fig. S4 and S2). Moreover, immunostaining of these PLA with LAMP2 confirmed that the interaction of p27 with Ragulator takes place on lysosomes (Fig. S4). The ability of p27 to interfere with Ragulator assembly was confirmed in MEFs, in which the endogenous LAMTOR5/LAMTOR1 interaction was lower in aa-starved p27^+/+^ MEFs compared to p27^−/−^ cells (Fig. 4J, K). Taken together, this data indicates that p27 binds to LAMTOR1 and interferes with Ragulator assembly (Fig. 4L).

An intact Ragulator complex is required for lysosomal targeting of Rag GTPases and depletion of LAMTOR1 or −2 results in cytoplasmic localization of both Rags and mTOR^23, 62^. Therefore, we tested if p27 impairs Rags recruitment to Ragulator. p27 expression in HEK293 cells decreased the amount of RagB immunoprecipitated with LAMTOR1 (Fig. 5A-C). Importantly, expression of a constitutively active Rag complex (RagB GTP/RagD GDP) was still affected by p27 expression (Fig. 5C), confirming that p27 acts upstream of Rag/Ragulator signaling, at the level of Ragulator assembly. Accordingly, the amount of endogenous LAMTOR4 co-precipitated with RagC was decreased in aa-starved p27^+/+^ MEFs, but it remained elevated in p27^−/−^ cells (Fig. 5D, E). Furthermore, the amount of RagA colocalizing with the lysosome marker LAMP2 was higher in aa-starved p27^−/−^ MEFs compared to p27^+/+^ cells, in which RagA appeared more diffuse in the cytoplasm, while they were similar under basal conditions (Fig. S5A, B). These results indicate that upon prolonged aa starvation, p27 prevents the assembly of Ragulator and the association of Ragulator with Rags on lysosomes.

**Figure 5:**
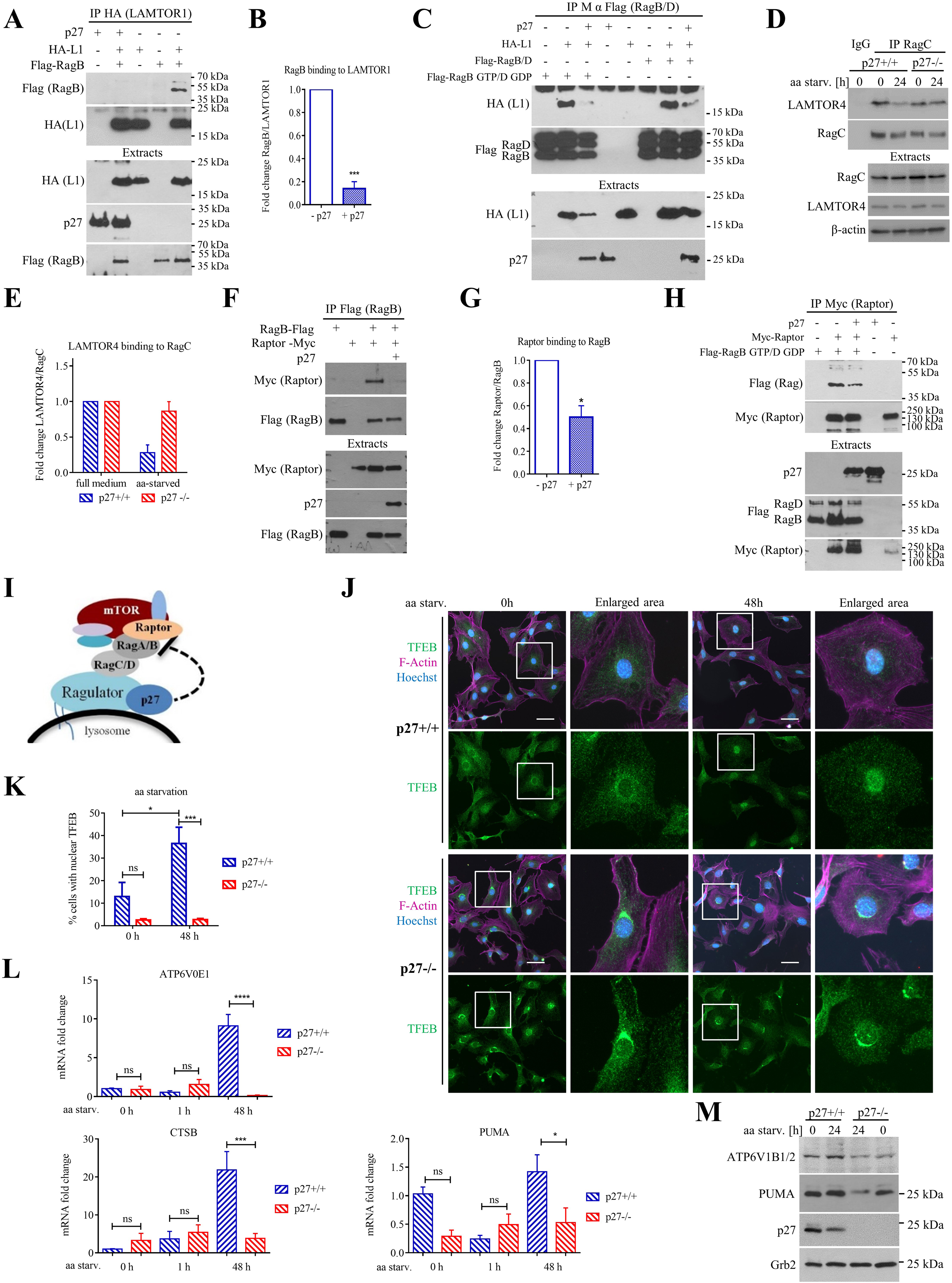
p27 interferes with Ragulator functions. **(A)** LAMTOR1 was immunoprecipitated with anti-HA antibodies from HEK293 cell expressing HA-LAMTOR1 and/or Flag-RagB and/or p27 and immunoblotted against Flag (RagB) and HA (LAMTOR1). Expression of the transfected proteins was verified by immunoblot of extracts with anti-Flag, −p27 and −HA antibodies. **(B)** Quantification of the amount of RagB co-precipitated with LAMTOR1 as in A from 5 experiments. Values were normalized to that in absence of p27. **(C)** RagB/D were immunoprecipitated with anti-Flag antibodies from HEK293 cell expressing HA-LAMTOR1 and/or Flag-RagB/D and/or Flag-RagB Q99L (RagB GTP)/RagD S77L (RagD GDP) and/or p27 and immunoblotted against Flag (RagB/D) and HA (LAMTOR1). Expression of transfected p27 and HA-LAMTOR1 in corresponding extracts are shown. **(D)** RagC was immunoprecipitated from p27^+/+^ and p27^−/−^ MEFs in full medium or aa starved for 24 h and the amount of LAMTOR4 co-precipitated was determined. Control IP with rabbit IgG was used as control. Levels of RagC and LAMTOR4 in the corresponding extracts are shown. β-actin was used as loading control. **(E)** Ratio of LAMTOR4 co-precipitated by that of RagC immunoprecipitated from p27^+/+^ and p27^−/−^ MEFs in full medium or aa starved for 24 h, normalized to full medium condition from 2 experiments as described in E. **(F)** RagB was immunoprecipitated with anti-Flag antibodies from HEK293 cells expressing Myc-Raptor and/or Flag-RagB and/or p27 and immunoblotted against Myc (Raptor) and Flag (RagB). Expression of the transfected proteins was verified by immunoblot of extracts with anti-Myc, −p27 and −Flag antibodies. **(G)** Quantification of the amount of Raptor co-precipitated with RagB as described in G from 3 experiments. Values were normalized to that in absence of p27. **(H)** Raptor was immunoprecipitated with anti-Myc antibodies from HEK293 cells expressing Myc-Raptor and/or Flag-RagB Q99L (RagB GTP)/RagD S77L (RagD GDP) and/or p27 and immunoblotted against Flag (RagB/D) and Myc (Raptor). Expression of transfected p27, Flag RagB/D and Myc-Raptor in corresponding extracts are shown. **(I)** Schematic summarizing the impact of p27 on Rag and mTORC1 recruitment to Ragulator. **(J)** TFEB immunostaining in p27^+/+^ and p27^−/−^ MEFs in full medium (0 h) or aa-starved for 48 h. F-actin was stained with phalloïdin and DNA with Hoechst. Scale bars are 50 µm. **(K)** Percentage of cells with nuclear TFEB signal in cells treated as in K from 3 experiments. At least 87 cells per condition for each genotype were analyzed in each experiment. **(L)** Fold change of the v-ATPase subunit ATP6V0E1, CTSB (Cathepsin-B) and PUMA mRNA levels in p27^+/+^ and p27^−/−^ MEFs in full medium or aa starved for 1 h or 48 h, determined by RT-qPCR and normalized to GAPDH levels from 8 experiments. All values were normalized to p27^+/+^ MEFs in full medium (0 h). **(M)** Immunoblot for the v-ATPase subunit ATP6V1B1/2, PUMA and p27 in p27^+/+^ and p27^−/−^ MEFs in full medium or aa starved for 24 h. Grb2 was used as loading control. **(B, E, G, K, L)** Bar graphs show means ± SEM. Statistical significance was evaluated by unpaired t-test with Welch’s correction (B, G) or 2-way ANOVA (K, L); ns: p >0.05; *: p ≤ 0.05; ***: p ≤ 0.001; ****: p ≤ 0.0001.

Rags recruit mTORC1 to lysosomal membranes by binding to the Raptor subunit of mTORC1^24^. Co-IP of RagB with Raptor was dramatically decreased when p27 was overexpressed in HEK293 cells, even when constitutively active RagB/D complex was used (Fig. 5F-H). Accordingly, mTOR immunostaining in p27^+/+^ MEFs became diffuse upon aa starvation, suggesting its release from lysosomal surface, while in p27^−/−^ MEFs mTOR colocalized with LAMP2 puncta even under aa starvation (Fig. S5C, D). Lysosomal localization of mTOR in p27^+/+^ MEFs was restored upon re-feeding cells with aa for 10 min (Fig. S5C, D). This data suggests that by interfering with Ragulator assembly, p27 inhibits the recruitment of Rags and mTOR to lysosomes in response to aa deprivation (Fig. 5I).

When nutrients are abundant, Rags recruit TFEB to lysosomes, where it is phosphorylated by mTOR, causing its cytoplasmic retention, while aa starvation leads to nuclear translocation of TFEB where it induces the transcription of pro-autophagy genes^29, 30, 75, 76^. Since p27 prevents Rag and mTOR recruitment to the lysosomal surface by interfering with Ragulator function, TFEB nuclear translocation in response to aa deprivation was used as another readout of Ragulator, Rag and mTOR activity in p27^+/+^ and p27^−/−^ MEFs. TFEB nuclear translocation was defective in aa-deprived p27^−/−^ cells, suggesting that mTOR remained active in these cells (Fig. 5J, K). Given that TFEB regulates a gene network that promotes lysosome biogenesis and autophagy^76^, impaired nuclear translocation of TFEB could underlie the autophagy defect in p27^−/−^ cells. Indeed, the expression of two TFEB target genes, *ATP6V0E1*, a v-ATPase subunit, and *CTSB*, encoding Cathepsin B, was dramatically reduced in aa-starved p27^−/−^ cells compared to p27^+/+^ MEFs (Fig. 5L), explaining the reduced levels of cathepsin B protein observed earlier (Fig. 3C). Moreover, expression of another v-ATPase subunit under TFEB control^76^, ATP6V1B2, was also reduced in aa-starved p27^−/−^ cells (Fig. 5M). Interestingly, expression of PUMA, another TFEB target involved in the induction of apoptosis^77^, was also strongly reduced in p27^−/−^ MEFs compared to p27^+/+^ cells (Fig 5L, M). Thus, several lines of evidence indicate that by binding to LAMTOR1, p27 interferes with Ragulator assembly and function, preventing the recruitment of Rags and mTOR to lysosomes and promoting TFEB nuclear translocation, thereby favoring autophagy.

### p27 participates in the inhibition of mTOR activity in amino acid deprived cells

Our data indicates that during aa starvation p27 inhibits mTOR recruitment to lysosomes and promotes TFEB nuclear translocation, suggesting that p27 may regulate mTOR activity. Indeed, phosphorylation of p70 S6K1 and 4E-BP1 at sites targeted by mTOR were elevated in p27^−/−^ cells aa starved for 48 h (Fig. 6A-C), consistent with the idea that p27 participates in mTOR inhibition. Furthermore, monitoring of mTOR phosphorylation itself by immunoblotting and immunostaining indicated that aa-starved p27^−/−^ cells maintained higher levels of active mTOR upon aa deprivation (Fig. 6D-G). Likewise, p27^CK−^ MEFs exhibited reduced P-p70 S6K1 levels, similar to p27^+/+^ cells, while p27^S10A^ cells maintained elevated P-p70 S6K1 levels, as in p27^−/−^ cells (Fig. 6H, I), indicating that the regulation mTOR activation is CDK-independent but requires p27 nuclear export. In line with these results, re-expression of p27 (Fig. 6J, K) or p27^CK−^ (Fig. 6L, M) in p27^−/−^ MEFs restored the inhibition of p70 S6K1 phosphorylation upon aa starvation, confirming that this effect is mediated by p27.

**Figure 6:**
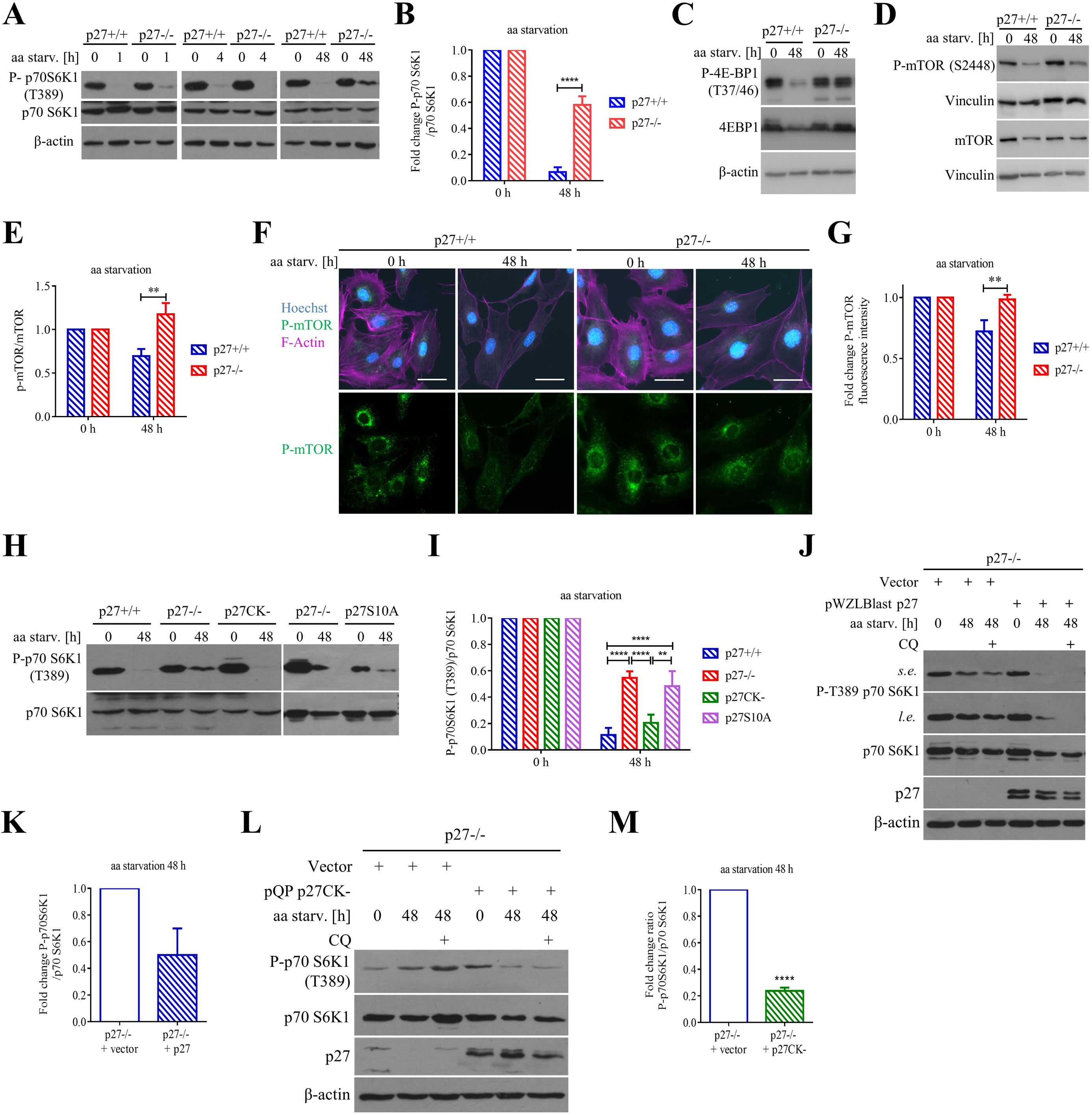
p27 participates in inhibition of mTOR signaling in amino acid deprived cells. **(A)** Immunoblots for P-T389 p70 S6K1 and p70 S6K1 in p27^+/+^ and p27^−/−^ MEFs in full medium or aa-deprived for the indicated times. β-actin was used as loading control. **(B)** Densitometry analysis of 5 experiments as described in A. P-p70 S6K1/p70 S6K1 ratio was normalized to control cells (0 h). **(C)** Immunoblots for P-T37/46 4E-BP1 and 4E-BP1 in p27^+/+^ and p27^−/−^ MEFs in full medium or aa-starved for 48 h. β-actin was used as loading control. **(D)** Immunoblot for P-S2448 mTOR and mTOR in p27^+/+^ and p27^−/−^ MEFs in full medium or aa-starved for 48 h. Vinculin was used as loading control. **(E)** Densitometry analysis of P-mTOR/mTOR ratio in 48 h aa-deprived cells normalized to cells in full medium from 3 experiments. **(F)** P-Ser2448 mTOR immunostaining in p27^+/+^ and p27^−/−^ MEFs in full medium or aa-starved for 48 h. F-actin was stained with phalloïdin and DNA with Hoechst. Scale bars are 50 µm. **(G)** Quantification of P-mTOR fluorescence intensity from 5 experiments as described in F, normalized to cells in full medium (0 h). **(H)** Immunoblot of P-T389 p70 S6K1 and p70 S6K1 in p27^+/+^, p27^−/−^, p27^CK−^ and p27^S10A^ MEFs in full medium (0 h) and aa starved for 48 h. **(I)** Densitometry analysis of P-p70 S6K1/p70 S6K1 ratio in 48 h aa-deprived cells normalized to cells in full medium from at least 3 experiments. **(J)** Immunoblot for P-T389 p70 S6K1 and p70 S6K1 in p27^−/−^ MEFs infected with empty or p27 expression vector in full medium or aa-starved for 48 h ± 50 µM CQ for 2 h. ‘s.e.’ = short exposure; ‘l.e.’ = long exposure. p27 levels are shown. β-actin was used as loading control. **(K)** Densitometry analysis of P-p70 S6K1/p70 S6K1 ratio in p27^−/−^ MEFs infected with empty or p27 expression vector aa-starved for 48 h from 2 experiments as described in J, normalized to empty vector infected cells. **(L)** Immunoblot of P-T389 p70 S6K1, p70 S6K1 and p27 in p27^−/−^ MEFs infected with empty or with p27^CK−^ expression vector in full medium (0 h) or aa-starved for 48 h ± 50 µM CQ for 2 h. p27 levels are shown. β-actin was used as loading control. **(M)** Densitometry analysis of P-p70 S6K1/p70 S6K1 ratio in p27^−/−^ MEFs infected with empty or p27^CK−^ expression vector aa-deprived for 48 h from 4 experiments as described in L, normalized to empty vector infected cells. **(B, E, G, I, K, M)** Bar graphs show means ± SEM. Statistical significance was analyzed by 2-way ANOVA (B, E, G, I) or unpaired t-test with Welch’s correlation (M). ****: p ≤ 0.0001; **: p ≤ 0.01.

Although cells respond to aa withdrawal by inhibiting mTOR activity and inducing autophagy, during prolonged aa starvation, autophagy-dependent replenishment of aa levels within lysosomes causes mTOR re-activation, which is required for ALR^24, 26, 32^. Importantly, mTOR activity is inhibited during short-time starvation in both p27^−/−^ and p27^+/+^ cells (Fig. 6A) and mTOR reactivation is detected in both genotypes, albeit with a marked increase in p27^−/−^ cells (Fig. 6A-G). To better understand how p27 regulates mTOR activity, the effect of blocking autophagy on mTOR reactivation was studied in aa starved cells. While autophagy inhibition with CQ blocked mTOR re-activation in p27^+/+^ MEFs, it had no effect in p27^−/−^ cells (Fig. 7A, B), suggesting that in absence of p27, mTOR signaling and autophagy are uncoupled. The requirement for autophagy to induce mTOR reactivation was confirmed in Tet-off Atg5^−/−^ MEFs in which Atg5 expression is turned off in presence of doxycycline, inhibiting autophagy^78^. In presence of doxycycline, mTOR reactivation was inhibited in these cells (Fig. 7C, D).

**Figure 7:**
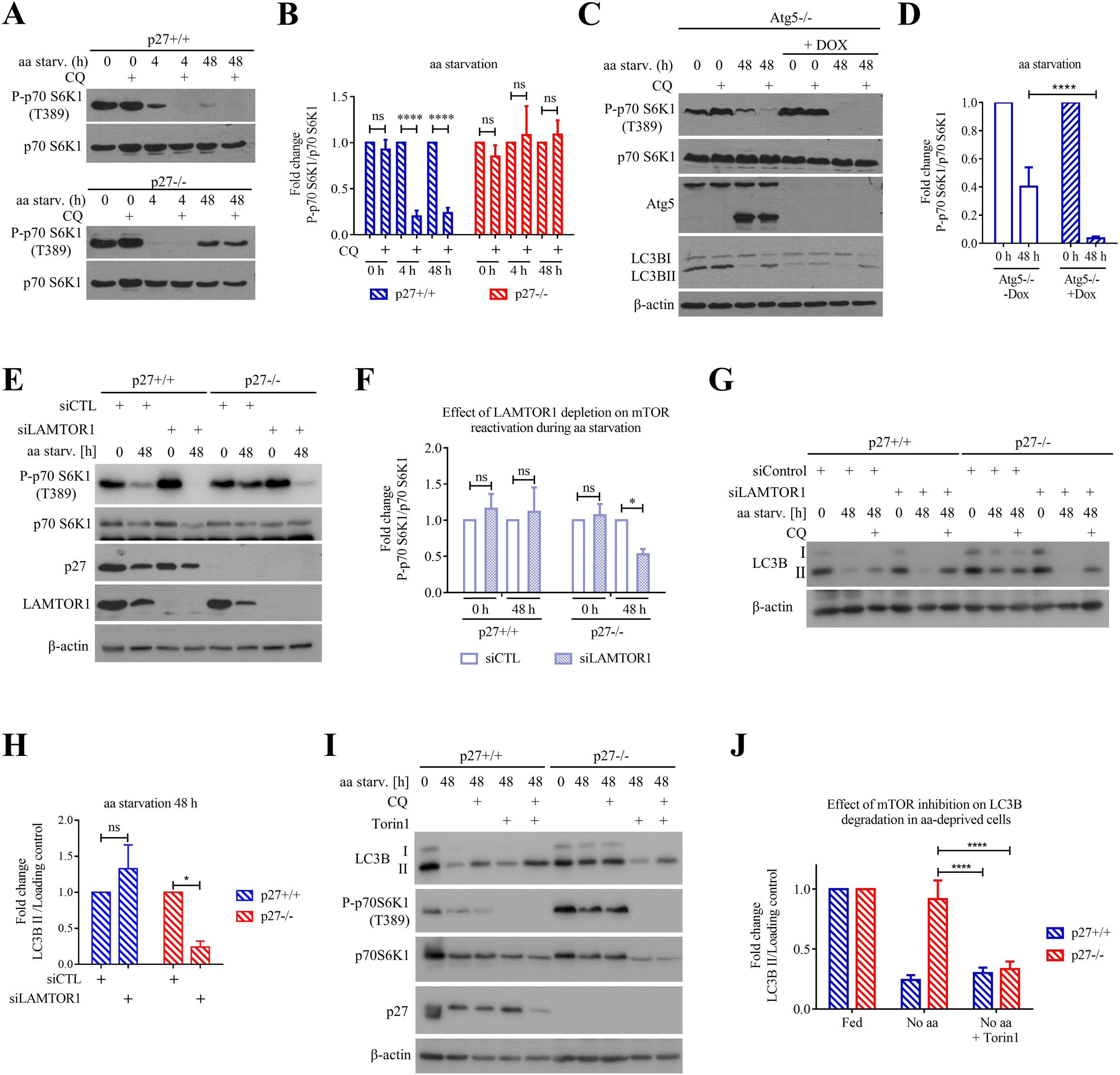
p27 controls mTOR and autophagy in a LAMTOR1 dependent manner. **(A)** Immunoblots for P-T389 p70 S6K1 and p70 S6K1 in p27^+/+^ and p27^−/−^ MEFs aa-starved for the indicated times ± 50 µM CQ for 2 h. mTOR reactivation is independent of autophagy in p27-null MEFs. **(B)** Densitometry analysis of P-p70 S6K1/p70 S6K1 ratio from 8 experiments as described in A. Values were normalized to condition without CQ for each time point. **(C)** Immunoblots for P-T389 p70 S6K1, p70 S6K1, LC3B and Atg5 in doxycycline-inducible Atg5^−/−^ MEFs treated or not with 10 ng/ml Doxycycline (Dox) in full medium (0 h) or aa deprived for 48 h ± 50 µM CQ for 2 h. β-actin was used as loading control. **(D)** Densitometry analysis of P-p70 S6K1/p70 S6K1 ratio from 4 experiments as described in C, normalized to cells in full medium. mTOR reactivation under prolonged aa starvation requires autophagy. **(E)** Immunoblots for P-T389 p70 S6K1, p70 S6K1, p27 and LAMTOR1 in p27^+/+^ and p27^−/−^ MEFs transfected with control or LAMTOR1 siRNA in full medium or aa-starved for 48 h. β-actin was used as loading control. **(F)** Quantification of P-p70 S6K1/p70 S6K1 ratio in cells from 8 experiments as described in E, normalized to control siRNA transfected cells in each condition. **(G)** Immunoblot for LC3B in p27^+/+^ and p27^−/−^ MEFs transfected with control or LAMTOR1 siRNA in full medium or aa starved for 48 h ± 50 µM CQ for 2 h. β-actin was used as loading control. **(H)** Quantification of LC3B-II/Loading control ratio from 5 experiments as described in G. Values were normalized to control siRNA transfected cells in each condition. **(I)** Immunoblots for LC3B, P-T389 p70 S6K1, p70 S6K1 and p27 in p27^+/+^ and p27^−/−^ MEFs in full medium or aa starved for 48 h ± 200 nM Torin1 for 24 h ± 50 µM CQ for 2 h. β-actin was used as loading control. **(J)** Densitometry analysis of LC3B-II/loading control ratio from 8 experiments as described in I, normalized to cells in full medium. **(B, D, F, H, J)** Bar graphs show means ± SEM. Statistical significance was evaluated by 2-way ANOVA (B, D, J) or multiple t test with Bonferroni correction (F, H); ****: p ≤ 0.0001; *: p ≤ 0.05; ns: p> 0.05.

To confirm that p27 controls mTOR activity and autophagy by interfering with Ragulator assembly and function, LAMTOR1 expression was silenced by siRNA in p27^−/−^ and p27^+/+^ cells (Fig. 7E). In full medium, LAMTOR1 silencing did not affect mTORC1 activation levels (Fig. 7E), consistent with Rag and Ragulator-independent pathways mediating mTORC1 activation, as reported previously^79, 80^. However, upon aa deprivation, LAMTOR1 knockdown in p27^−/−^ MEFs restored mTOR inhibition, evaluated by measuring p70 S6K1 phosphorylation levels (Fig. 7E, F), and LC3B degradation (Fig. 7G, H), without having a significant effect in p27^+/+^ cells, indicating that p27 regulates mTOR activation state and autophagy via a LAMTOR1-dependent mechanism. Similarly, mTOR inhibition with Torin1, confirmed by P-p70 S6K1 immunoblot (Fig. 7I), restored LC3B-II degradation in aa starved p27^−/−^ cells (Fig. 7I, J), confirming that p27 controls autophagy via an mTOR-dependent mechanism. Taken together, this data indicate that p27 participates in the inhibition of mTOR activity during prolonged aa deprivation and this is important for autophagy induction. Conversely, p27^−/−^ cells maintain elevated mTOR activity and exhibit reduced autophagy flux.

### Elevated mTOR activity in p27^−/−^ cells confers resistance to amino acid starvation-induced apoptosis

p27 was previously found to promote survival under serum and glucose starvation^14, 36, 37^. Surprisingly, p27 had opposite effects on survival upon glucose or aa starvation. While p27 expression promoted survival in glucose deprived MEFs, consistent with previous reports^14, 36, 37^, aa starved p27^+/+^ MEFs were markedly more susceptible to apoptosis than p27^−/−^ cells (Fig. 8A, B). The specificity of this approach to measure apoptosis was validated by addition of ZVAD pan-caspase inhibitor, which blocked apoptosis in aa deprived cells (Fig. S6A, B). Differences in caspase cleavage were confirmed by immunoblot (Fig. S6C). Re-expression of p27 in p27^−/−^ MEFs increased their susceptibility to apoptosis upon aa starvation (Fig. S6D, E), confirming the implication of p27 in this phenotype.

**Figure 8:**
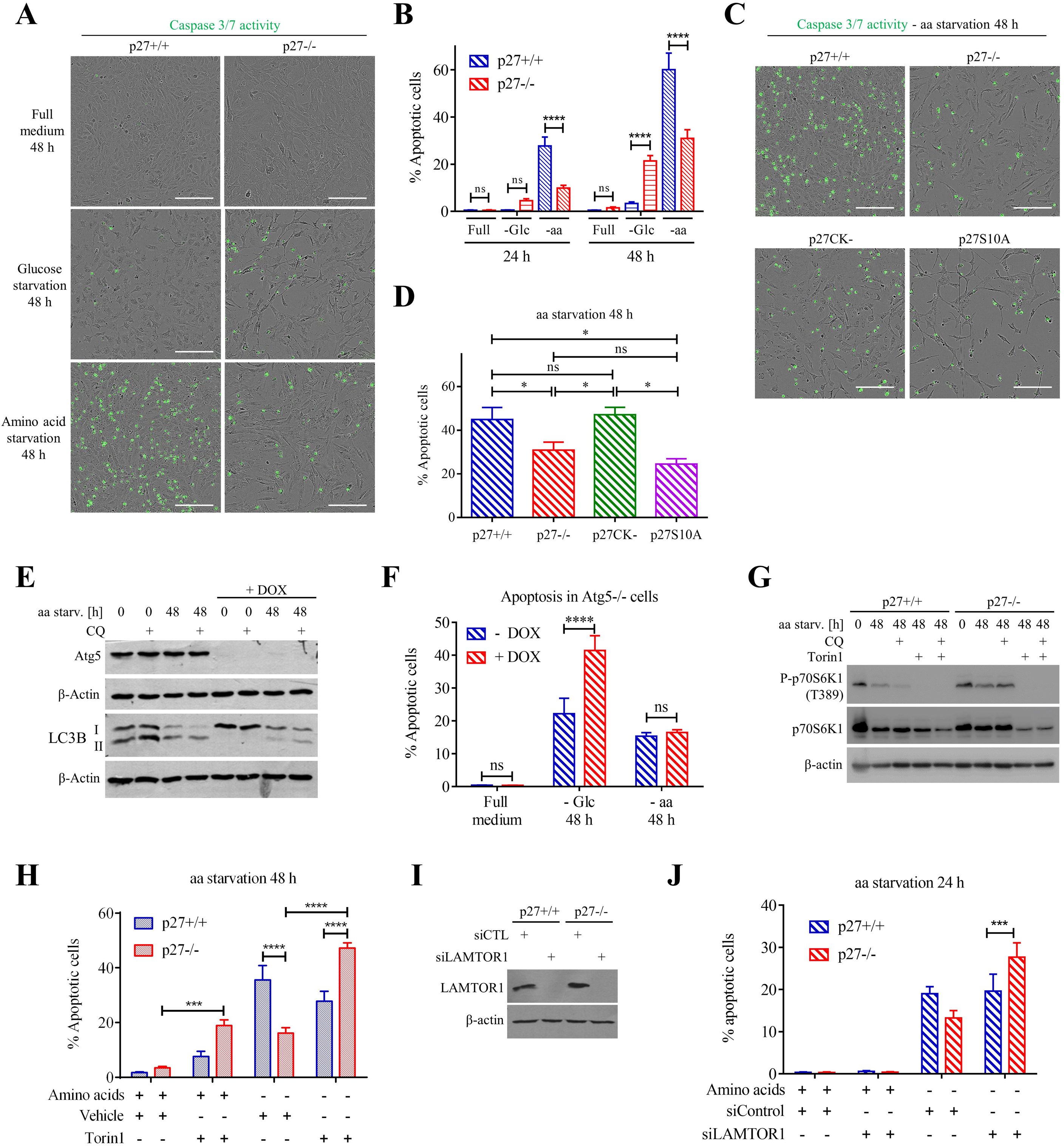
Elevated mTOR activity in p27^−/−^ cells confers resistance to amino acid starvation-induced apoptosis. **(A)** p27 status determines cell survival in response to metabolic stress. Incucyte images of phase contrast and green fluorescence representing caspase-3/7 cleavage of p27^+/+^ and p27^−/−^ MEFs grown for 48 h in full medium or in absence of glucose or aa. Scale bars are 200 µm. **(B)** Percentage of apoptosis in p27^+/+^ and p27^−/−^ MEFs in full medium or glucose or aa starved for 24 h and 48 h from at least nine experiments. **(C)** Incucyte images of p27^+/+^, p27^−/−^, p27^CK−^ and p27^S10A^ MEFs aa-starved for 48 h. Scale bars are 200 µm. **(D)** Percentage of apoptosis in MEFs treated as in C from at least 3 experiments. **(E)** Immunoblot for ATG5 and LC3B in ATG5 inducible knockout MEFs ± dox in full medium or aa starved for 48 h ± 50 µM CQ for 2 h. **(F)** Percentage of apoptotic ATG5 inducible knockout MEFs ± dox in full medium or glucose or aa starved for 48 h from at least 15 experiments. Images for these experiments are shown Fig. S6F. **(G)** Immunoblotting of P-p70 S6K1, p70 S6K1 and p27 in p27^+/+^ and p27^−/−^ MEFs in full medium or aa-starved for 48 h ± 200 nM Torin1 ± 50 µM CQ for 2 h. β-actin was used as loading control. **(H)** Percentage of apoptotic p27^+/+^ and p27^−/−^ MEFs in full medium or aa-starved for 48 h ± 200 nM Torin1 from eight experiments. Images for these experiments are shown Fig. S6G. **(I)** Immunoblot for LAMTOR1 and β-actin of p27^+/+^ and p27^−/−^ MEFs transfected with control or LAMTOR1 siRNA. **(J)** Percentage of apoptotic p27^+/+^ and p27^−/−^ MEFs in full medium or aa-starved for 48 h transfected with control or LAMTOR1 siRNA from ten experiments. Images for these experiments are shown Fig S6H. **(B, D, F, H, J)** Bar graphs show means ± SEM. Statistical significance was evaluated by 2-way ANOVA (B, F, H, J) or one-way ANOVA (D) tests; ns: p > 0.05; *: p ≤ 0.05. ***: p ≤ 0.001; ****: p ≤ 0.0001; *****: p ≤ 0.00001.

The role of p27 in protecting against starvation-induced apoptosis was associated with its cytoplasmic localization and capacity to inhibit CDK activity^14, 34, 36^. The importance of these features were tested in aa deprivation conditions. p27^S10A^ MEFs were resistant to aa deprivation-induced apoptosis, similar to p27^−/−^ cells, indicating that the cytoplasmic localization of p27 is required to promote apoptosis (Fig. 8C, D). On the other hand, p27^CK−^ MEFs behaved like p27^+/+^ cells and were highly susceptible to aa deprivation-induced apoptosis, indicating that the regulation of cyclin-CDK complexes is not involved in this process (Fig. 8C, D). This was confirmed by re-expression of p27^CK−^ in p27^−/−^ MEFs, which restored the susceptibility to aa deprivation-induced apoptosis similar to p27^+/+^ cells (Fig. S6D, E). Thus, while p27 expression promotes survival in response to glucose starvation, it plays a pro-apoptotic role during aa deprivation and this pro-apoptotic role is CDK-independent but requires p27 nuclear export.

Cells lacking p27 exhibit a survival advantage during prolonged aa starvation conditions, but also display elevated mTOR activity and decreased ability to perform autophagy. First, the importance of autophagy in mediating survival upon glucose or aa-starvation was tested using Tet-off Atg5^−/−^ MEFs. Loss of Atg5 impaired autophagy (Fig. 8E), as expected^78^, and dramatically increased glucose starvation-induced apoptosis, but had no effect on survival upon aa deprivation (Fig. 8F and S6F). Thus, autophagy promotes survival in absence of glucose, and reduced autophagy in p27^−/−^ cells most likely underlies their susceptibility to glucose starvation-induced apoptosis. Second, we tested whether the elevated levels of mTOR activity in p27^−/−^ MEFs were responsible for promoting survival, as mTOR is known to regulate cell survival in a context-specific manner^81, 82^. mTOR inhibition with Torin1 (Fig. 8G) did not increase cell death in aa-starved p27^+/+^ MEFs, in which mTOR activity is already low, but potently caused apoptosis in p27^−/−^ cells (Fig. 8H and S6G). LAMTOR1 silencing also reversed the resistance to aa starvation-induced apoptosis of p27^−/−^ MEFs (Fig. 8I, J and S6H), confirming the importance of p27-mediated regulation of Ragulator assembly and function in this process. Taken together, this data indicates that the resistance of p27^−/−^ cells to prolonged aa starvation-induced apoptosis is not a consequence of impaired autophagy, but of their ability to maintain mTOR activity and signaling, notably preventing TFEB activation and suppressing expression of the pro-apoptotic protein PUMA.

## DISCUSSION

In all living organisms, proliferation and growth must be tightly coordinated during development and to maintain homeostasis. The prevailing view, from experiments in yeast, flies and mammals is that growth signals are dominant over cell cycle control^1^. Indeed, when an organism or organ reaches its predetermined size or following metabolic restriction, growth and proliferation coordinately cease. The cyclin/CDK inhibitor p27 plays a major role in the regulation of cell division by causing cell cycle arrest in G1 phase^2, 83^. This role is underscored by the phenotype of p27 knockout mice that exhibit gigantism, with an overall increase in body size of approximately 30%^4^. In these animals, organs grow beyond their normal size and cells fail to enter quiescence in a timely manner, possibly because they are refractory to growth inhibitory signals. mTOR is a master regulator of growth and its activation or inhibition dictates whether cells adopt a catabolic or anabolic metabolism, respectively^25^. p27 has long been known as a major effector of cell cycle arrest following mTOR inhibition by rapamycin, which induces p27 expression at the transcriptional and post-transcriptional levels^84–87^. Conversely, mTOR activity drives p27 levels down, notably by inducing Myc, cyclin E, cyclin D and Skp2 expression^86, 88^, and causes p27 cytoplasmic localization by activating the kinase SGK1, which phosphorylates p27 on T157^89^. Here, we find that prolonged aa starvation causes the relocalization of a fraction of p27 to lysosomal membranes, where it interacts with LAMTOR1, preventing Ragulator assembly and participating in mTOR inhibition. Thus, it appears that upon metabolic stress, p27 can directly exert a negative feedback on mTOR signaling, providing an example of cross talk between the cell cycle and cell growth machineries. In these conditions, p27 acts both as a cell cycle inhibitor and a growth inhibitor, preventing anabolic activity from restarting before metabolic conditions have sufficiently improved, therefore coordinating growth and proliferation. Recent evidence indicate that other components of the cell cycle machinery are also involved in metabolic control at the transcriptional level. The CDK4/Rb/E2F1 pathway drives the expression of a number of genes involved in metabolism, notably in mitochondrial function and lipogenesis, possibly allowing metabolic adaptation to the proliferative state of specific tissues^90, 91^.

Our data provides mechanistic insight in the role of p27 in autophagy in response to aa deprivation. While p27 is known to play a pro-autophagic role under basal conditions, serum and glucose starvation^14, 33, 34, 37^, its effect on aa deprived cells was completely unknown. Interestingly, during prolonged aa starvation, the regulation of mTOR activity required the cytoplasmic localization of p27 but was independent of its ability to regulate CDKs, as p27^S10A^ and p27^CK−^ MEFs behaved like p27^−/−^ and wild type cells, respectively. In contrast, during glucose starvation, CDK inhibition by p27 was required to control autophagy^14, 34^. Furthermore, AMPK signaling, which played a major role in p27-mediated autophagy in glucose or serum starved cells^14^, was not activated in aa deprived cells, possibly due to the presence of glucose and serum in the starvation medium, and p27 phosphorylation on T198 did not increase in our experiments. Thus, it appears that p27 regulates autophagy by distinct mechanisms in response to different metabolic stresses.

During prolonged aa starvation, p27^−/−^ cells exhibit increased mTOR activity and TFEB cytoplasmic sequestration, which decreases lysosomal function and autophagy flux, leading to accumulation of autophagic cargo. Interestingly, autophagy induction initially occurs normally in both wild type and p27^−/−^ cells and it is only upon prolonged starvation that p27^−/−^ cells display enhanced mTOR reactivation and impaired autophagy. This correlates with the recruitment of p27 to autophagic compartments. Localization of p27 in autophagic compartment was previously suggested but its functional significance was unclear^14, 92, 93^. An attractive hypothesis is that p27-mediated inhibition of mTOR intervenes only during sustained metabolic stress to enforce inhibition of growth promoting signals.

Regulation of autophagy is a complex process in which mTOR activity must be tightly coordinated to control all stages of autophagy, from autophagosome formation to recycling of lysosomes during prolonged periods of starvation. We found that cytoplasmic p27 plays a crucial role in the regulation of autophagy during prolonged aa starvation by controlling mTOR re-activation via the modulation of Ragulator activity. Although mTOR inhibits early autophagy events by phosphorylating ULK1, preventing autophagosome formation, ULK1 levels were very low following prolonged aa starvation, as previously reported^47^, suggesting an ULK1-independent mechanism in these conditions. The suppression of mTOR activity and nuclear translocation of TFEB are required for activation of lysosomal functions and cargo degradation during autophagy^94^. Our data suggests that in aa-deprived p27^−/−^ cells, enhanced mTOR activity partially prevents TFEB activation by inhibiting its nuclear translocation and the subsequent activation of lysosome-related genes, including v-ATPase, which is required to maintain a low pH in the lysosomal lumen^95^, and Cathepsin-B, and this is associated with decreased acidification and impaired lysosomal proteolysis^76^.

Lysosome-anchored Ragulator recruits Rag GTPase heterodimers, and when Rags are properly loaded (RagA or B GTP and RagC or D GDP), they capture the Raptor subunit of mTORC1 to lysosomal membranes^96^. There, mTOR can be activated by Rheb, itself under control of the PI-3K/AKT/TSC signaling pathway. This complex mechanism of regulation ensures a tight control of mTOR activity. Our data suggests that binding of p27 to LAMTOR1 interferes with the interaction of LAMTOR1 with three other subunits of the Ragulator complex, LAMTOR2, −3 and −5, but not with LAMTOR4, preventing Ragulator assembly and Rag recruitment. This is supported by the fact that p27 also prevents constitutively active Rag GTPase complexes from binding to Ragulator or to Raptor. Interestingly, solving of the structure of Ragulator in complex with Rag GTPases shows that LAMTOR1 adopts a horseshoe shape that accommodates the other LAMTOR subunits, with LAMTOR4 sitting at the bottom of the horseshoe and Rags associating with the tips of LAMTOR1 U-shape^72–74^. This is consistent with p27 straddling LAMTOR1 and preventing the interaction with the partners that bind to the distal ends of LAMTOR1 (i.e. LAMTOR2, −3 and −5). Structural studies on the p27/LAMTOR1 interaction and Ragulator assembly will be very informative to further understand how p27 affects Ragulator function.

Surprisingly, despite reduced autophagy levels, cells lacking p27 were partially resistant to aa starvation-induced apoptosis, unlike what was previously reported under serum and glucose starvation^14, 36, 37^. In fact, we found that autophagy did not protect cells in this context, as pharmacological or genetic inhibition of autophagy did not affect survival under aa starvation. Instead, elevated mTOR activity in p27^−/−^ appeared to be responsible for increased survival, since it could be reversed by pharmacological inhibition of mTOR or LAMTOR1 silencing. An attractive hypothesis is that active mTOR inhibits TFEB activity by cytoplasmic retention, which prevents the induction of pro-apoptotic PUMA, as recently reported^77^. Elevated mTOR activity could also promote survival via the phosphorylation of Bad on S136, inactivating its pro-apoptotic function^97^. Alternatively, mTOR inhibition was recently found to reduce lysosomal efflux of most essential amino acids^98^, suggesting that p27 expressing cells, which exhibit low mTOR activity, could have a reduced ability to reuse aa compared to p27^−/−^ cells following aa deprivation-induced autophagy, leading to cell death. p27 status appears to be a critical determinant in the ability of cells to survive to different metabolic stress: while p27-null cells survive to aa-starvation, they are more sensitive to glucose deprivation. These findings may be particularly relevant in the context of cancer, in which targeting specific metabolic pathways in function of p27 status, i.e. glucose metabolism in p27-low cells and aa metabolism in p27-high cells could improve treatment response. Moreover, our data implies that using mTOR inhibitors in conjunction with aa metabolism targeting drugs could overcome the resistance to targeting aa metabolism alone. Interestingly, p27 was identified as a potential predictive biomarker to the response to mTOR inhibitors^99^. Our results certainly warrant further investigations to test these possibilities.

Overall, in prolonged aa deprivation conditions, p27 can impinge on mTOR signaling by inhibiting Ragulator assembly, thereby promoting autophagy. These results provide a direct link between cell cycle control and growth signaling.

## MATERIALS AND METHODS

### Antibodies, reagents and plasmids

Mouse anti p27 (F8, sc-1641), p27 (SX53G8.5, sc-53871), Flag (H5, sc-166355), HA (F7, sc-7392), LAMP1 (E5, sc-17768), Myc (9E10, sc-40), 4E-BP1 (R113, sc-6936), PUMA (G3, sc-374223) and rabbit anti p27 (C19, sc-528), Myc (A14, sc-789), HA (Y11, sc-805), Flag (D8, sc-807) antibodies were purchased from Santa Cruz Biotechnology. Mouse anti p27 (610242) and Grb2 (610112) antibodies were purchased from BD-Transduction Laboratories. Mouse anti β-actin (A2228), Vinculin (V9131) and β-tubulin (T4026) antibodies were purchased from Sigma-Aldrich. Rabbit anti phospho-T198 p27 was purchased from R&D Systems. Rabbit anti phospho-p70 S6K1 (Thr389) (#9234), p70 S6K1 (#27087), mTOR (#2983), phospho-mTOR (Ser2448) (#5536), Raptor (#2280), LAMTOR1 (#8975), LAMTOR4 (#13140), LAMTOR5 (#14633), LC3B (#4108), RagA (#4357), RagC (#5466), Cathepsin-B (#D1C7Y), ATP6V1B1/2 (#13569), ULK1 (#8054), phospho-ULK1 (Ser758) (#6888), AMPK (#5832), phospho-AMPK (Thr172) (#2535), p62/SQSTM1 (#39749), phospho-4E-BP1 (Thr37/46) (#2855) and Atg5 (#12994) antibodies were purchased from Cell Signalling Technology. Rabbit anti TFEB (A303-673A-T) antibody was purchased from Bethyl Laboratories. Rat anti LAMP2 (Ab13524) antibody was purchased from Abcam. Mouse anti LC3B (0231-100/LC3-5F10) was purchased from NanoTools. Secondary antibodies against whole Ig or Ig light-chain conjugated to horseradish peroxydase or Cyanine-2 and −3 were purchased from Jackson ImmunoResearch. Phalloidin-Fluoprobes647 (FP-BA0320) was purchased from Interchim. Control (sc-108727) and Mouse LAMTOR1 (sc-37007) siRNA were purchased from Santa Cruz Biotechnologies. Lysotracker Deep Red (L12492) was purchased from Thermo Fisher. Self-Quenched BODIPY FL conjugate of BSA (Green) (#7932) was purchased from BioVision. Tetramethylrhodamine isothiocyanate– Dextran (T1037), Chloroquine diphosphate (C6628) and HRP-conjugated Protein G (#18-160) were purchased from Sigma-Aldrich. Torin1 (#4247) was purchased from Tocris. Recombinant His-tagged LAMTOR1 (#CSB-EP757083HU) was purchased from Cusabio Technology LLC.

p27 constructs and p27 point mutants and deletion mutants in pCS2+, pcDNA3.1+Hygro (Invitrogen), pQCXIP (Clontech), pBabe-puro, pWZL-Blast and pGEX4T1 were described previously^10, 12, 13^.

pBabe-puro-mCherry-eGFP-LC3B was a gift from Jayanta Debnath (Addgene #22418)^100^. pRK5 Flag-p18 (LAMTOR1) (Addgene #42331), pRK5 HA-p18 (LAMTOR1) (Addgene #42338), pRK5 HA C7orf59 (LAMTOR4) (Addgene #42336), pRK5 Flag C7orf59 (LAMTOR4) (Addgene #42332), pRK5 Flag p14 (LAMTOR2) (Addgene #42330), pRK5 HA mp1 (LAMTOR3) (Addgene #42329), pRK5 HA HBXIP (LAMTOR5) (Addgene #42328) and pRK5 Flag HBXIP (LAMTOR5) (Addgene #42326)^62^, pRK5 Myc-Raptor (Addgene #1859)^101^, and pLJM1 Flag-RagB (Addgene #19313), pLJM1 Flag-RagB Q99L (Addgene #19315), pLJM1 Flag-RagD (Addgene #19316), pLJM1 Flag-RagD S77L (Addgene #19317)^24^ were gifts from David Sabatini. All plasmids were verified by DNA sequencing.

### Cell culture and Transfections

Primary MEFs were prepared as described previously from p27^+/+^, p27^CK−/CK−^, p27^S10A/S10A^ or p27^−/−^ embryos^6, 13, 102^. MEFs were immortalized by infection with retroviruses encoding the human papilloma virus E6 protein and hygromycin selection. Retroviral infections were performed as described previously^13^. The following concentrations of antibiotics were used for selection: 2 µg/mL puromycin, 250 µg/mL hygromycin, 16 µg/mL blasticidin. Cells were kept under selection at all times.

All cells were grown at 37°C and 5% CO_2_ in DMEM (D6429, Sigma), 4.5 g/l glucose supplemented with 10% fetal bovine serum [FBS], 0.1 mM nonessential amino acids and 2 µg/ml penicillin-streptomycin. For starvation experiments, cells were rinsed twice with PBS and once with aa starvation medium (DMEM low glucose no amino acids [D9800, USBiological] complemented with glucose to 4.5 g/l, 0.1 mM sodium pyruvate, 2 µg /ml penicillin-streptomycin and 10% dialyzed FBS) and kept in starvation medium for the indicated times. For re-addition experiments, starvation medium was replaced with aa containing medium (D6429 Sigma) for the indicated time. For all starvation experiments, FBS was dialyzed against PBS in 3,500 MW cut-off dialysis tubing (SpectrumLabs, 132111) following manufacturer’s instructions.

Where indicated, 50 µM Chloroquine for 2 h and/or 200 nM Torin1 for 24 h (immunoblot)s or 48 h (apoptosis assays), or 20 µM ZVAD for 48 h, were added to the medium. Control cells were treated with identical volume of vehicle. For experiments using Tet-off inducible Atg5^−/−^ MEFs, kindly provided by Dr. N. Mizushima (Metropolitan Institute of Medical Science, Tokyo, Japan)^78^, cells were grown for 4 days in medium containing 10 ng/ml doxycycline prior to and during starvation. For Lysotracker staining, cells were incubated with 100 nM of Lysotracker Deep Red (ThermoFisher, L12492) in normal or aa starvation medium for 1 h. Cells were washed with PBS before microscopy analysis. Lysotracker fluorescence intensity was measured with the Nikon NIS Element software.

HEK293 and U251N cells were authenticated by short tandem repeat profiling. All cells were routinely tested to be free of mycoplasma contamination by DAPI staining. HEK293 cells were transfected by the calcium phosphate method 24 h prior to lysis. siRNA transfection was performed using Interferin (Polyplus transfection) according to manufacturer’s instructions 48 h before starvation.

### Immunoprecipitation and GST Pull-down

Cells were scraped and lysed in IP buffer (1% NP-40, 50 mM HEPES pH 7.5, 1 mM EDTA, 2.5 mM EGTA, 150 mM NaCl, 0.1% Tween 20 and 10% glycerol, complemented with 1 mM DTT, phosphatase inhibitors (10 mM β-glycerophosphate, NaF, sodium orthovanadate) and protease inhibitors (10 µg/ml Aprotinin, Bestatin, Leupeptin and Pepstatin A). For LAMTOR1 IPs, NP40 was replaced with 1% ODG (Octyl-β-D-glucopyranoside, O8001, Sigma) in the lysis buffer. After sonication for 10 sec, cells extracts were centrifuged for 5 min at 12,500 rpm and supernatants were collected. Protein concentration was quantified using Bradford reagent (BioRad). Lysates (500 µg for HEK293 or 2 mg for U251N) were incubated with 3 µg of indicated antibodies and 12 µl protein-A sepharose beads (IPA300, Repligen) (co-IP) or with recombinant GST proteins and glutathione sepharose beads (Pharmacia) (GST pull-down) at 4°C for 4 h. Beads were then washed 4 times in lysis buffer and resuspended in 10 µl 4X Laemmli buffer, boiled 3 min at 96°C, and subjected to immunoblot.

### Immunoblot

Cells were lysed either in IP buffer as described above or directly in 2X Laemmli buffer. Lysates and immunoprecipitates were mixed with 4X Laemmli buffer and boiled. Proteins were resolved on 8-15% SDS-PAGE depending on protein size and transferred to polyvinylidene difluoride membrane (Immobilon-P, Millipore). Membranes were blocked with PBS-T (PBS, 0.1% Tween-20), 5% non-fat dry milk and probed with indicated primary antibodies overnight at 4°C with agitation. Membranes were washed 3 times in PBS-T prior to incubation with corresponding HRP-conjugated secondary antibody (1/10000) or Protein G-HRP for LAMTOR1 IPs for 4-6 h at room temperature. Bands were visualized using enhanced chemiluminescence detection reagents (Millipore, BioRad, and Ozyme) and autoradiographic films (Blue Devil) or with a Fusion Solo S (Vilber) digital acquisition system.

To monitor endogenous LAMTOR1 levels, cells were lysed in 2X Laemmli buffer (4% SDS, 20% Glycerol, 120 mM Tris-HCl pH 6.8). Lysates were subjected to two 15 sec rounds of sonication. Then cells were centrifuged for 5 min at 12,500 rpm and supernatants were collected. Bromophenol blue 0.02% and DTT at final concentration 200 mM were added after BCA quantification. Lysates were boiled for 3 min at 96°C prior to electrophoresis.

Intensity of Western Blot signal were evaluated by densitometry analysis using the ImageJ software and normalized to loading control (β-Actin, β-Tubulin or Grb2) density value. Phospho-protein signals were normalized to the corresponding total protein levels. For LC3B turnover assays, LC3B/loading control ratio was measured in presence and absence of CQ and the ratio of these values was interpreted as a rate of autophagy flux. For co-IP quantifications, co-precipitated protein was normalized to precipitated protein in the same condition.

### Lysosome purification

Lysosomal enrichment was performed using the Lysosome Isolation Kit LYSIO1 (Sigma-Aldrich). Cells were trypsinized and suspended in 1X Extraction buffer complemented with 1 % protease inhibitors prior to homogenization in dounce with a B pestle. Resulting extracts were centrifuged at 1000 g for 10 min at 4°C. The supernatant was collected and centrifuged at 20,000 g for 20 min at 4°C. The resulting supernatant was collected as ‘cytoplasm’ fraction. The pellet was resuspended using a pellet pestle in 1X Extraction buffer to obtain the Crude Lysosomal Fraction (CLF). The CLF was centrifuged at 150,000 g at 4°C for 4 h to remove mitochondria and ER ^51^. The final pellet constituted the ‘lysosome’ fraction and was resuspended in 1X Laemmli buffer and sonicated for 10 s. BCA protein Assay Kit (Sigma-Aldrich) was used to quantify proteins in all the fractions and 30 µg of proteins per fraction were separated by SDS-PAGE for immunoblot.

### BSA dequenching assay

Cells were seeded on glass coverslips until 60-80% confluence and aa starved for 48 h. Self-Quenched BODIPY FL Conjugate of BSA (DQ-BSA) was added at 10 µg/mL 1 h before the end of experiment. Coverslips were rinsed three times in PBS and fixed with 2% PFA for 20 min at 37°C prior to microscopy analysis.

### Dextran labeling

Cells were seeded on glass coverslips and incubated in dextran-containing medium (20 mg/mL) for 18 h. Cells were then washed 3 times with dextran-free medium and incubated with dextran-free medium for 3 h. Coverslips were rinsed three times with PBS and fixed with 2% PFA for 20 min at 37°C prior to fluorescence microcopy.

### Immunofluorescence

Cells were seeded on coverslips and grown to 80-90% confluence before proceeding to starvation for the indicated times. Cells were rinsed with PBS and fixed with either 2% PFA in PBS for 20 min at 37°C or with 1% PFA for 3 min at room temperature followed by 100% methanol for 5 min at −20°C. For immunostaining, cells were permeabilized for 3 min with PBS 0.2% Triton X-100, except for LAMP2 staining that required permeabilization with 0.1% saponin in PBS, rinsed three times 5 min in PBS and incubated for 20 min in blocking solution (PBS, 3% BSA, 0.05% Tween 20 and 0.08% sodium azide) and with primary antibodies diluted in blocking solution for 1 h at 37°C. After three 5 min washes in PBS, cells were incubated for 30 min at 37°C with Cy2, Cy3 or Cy5-conjugated secondary antibodies at 1/500 dilution. In some experiments phalloidin-Fluoprobes 647 at 1/500 was added to secondary antibody solution. Next, coverslips were washed 3 times 5 min in PBS, with the first wash containing 0.1 μg/ml Hoechst H33342. Coverslips were mounted on glass slides with gelvatol (20% glycerol (v/v), 10% polyvinyl alcohol (w/v), 70 mM Tris pH 8). Images were captured on a Nikon 90i Eclipse microscope using a Nikon DS-Qi2 HQ camera. NIS Element BR software was used for acquisition and image analysis. For co-localizations, fluorescence intensity profiles were generated for each channel using NIS Element software. Overlapping of peaks between two channels was considered as a double-positive area. To measure fluorescent intensity of specific cellular compartments, regions of interest (ROI) were delineated and the signal was analyzed within the ROI.

### Proximity Ligation Assay

PLA was performed using Duolink *in situ* fluorescence technology (Sigma-Aldrich) according to the manufacturer’s protocol. Briefly, cells were plated on glass coverslips and grown overnight before starvation. Cells were fixed in 2% formaldehyde for 20 min at room temperature and permeabilized with either 0.2% Triton X-100 or 0.1% saponin in PBS for 3 min. Cells were blocked with Duolink blocking solution for 30 min at 37°C and incubated with combinations of primary antibodies for 1 h at 37° C. The following antibodies were used in PLA assays: mouse anti p27 (SX53G8.5, sc-53871), LAMP1 (E-5, Sc-17768), rabbit anti p27 (C19, sc-528) (Santa Cruz Biotechnology), rabbit anti LAMTOR1 (#8975), LAMTOR4 (#13140) (Cell Signalling Technology) at 1/200 dilution. Coverslips were incubated with secondary antibodies, conjugated with PLA probes anti-Rabbit PLUS (DUO92002) and anti-Mouse MINUS (DUO92004) for 1 hour at 37°C. Duolink PLA Detection Reagent Red (DUO9008) was used according to manufacturer’s instructions. After the amplification step, cells were incubated with coupled Phalloidin-A488 at 1/500 for 30 min. To visualize lysosomes, after the PLA, coverslips were incubated with rat anti LAMP2 (GL2A7, ab13524) at 1/400 dilution, followed by secondary antibody at 1/500. DNA was stained with 0.1 μg/ml Hoechst 33342. Images were captured on a Nikon 90i Eclipse microscope using a Nikon DS-Qi2 HQ camera. PLA dots and nuclei were counted using the Image J software.

### RNA extraction and RT-qPCR

Cells were lysed in TRI reagent (T9424, Sigma) and RNA was isolated according to manufacturer’s protocol. The integrity of RNA was verified on 0.8% agarose gel and quantified on NandoDrop. cDNA was synthesized used using SuperScipt III or SuperScirpt IV (ThermoFisher) according to manufacturer’s instructions using 1 µg of template RNA per reaction.

qPCR was performed in Bio-Rad CFX96 plates using SsoFast EvaGreen Supermix (Bio-Rad) using primers at a final concentration of 500 nM and an amount of cDNA corresponding to 2,5 ng of RNA per reaction. Bio-Rad CFX96 Real-Time PCR system was used to generate Ct values. Data was analyzed and normalized by the 2^−ΔΔ*CT*^ method using GAPDH as a housekeeping gene. All Ct values were normalized to gene expression in untreated p27^+/+^ cells in the same experiment.

Primers used were:

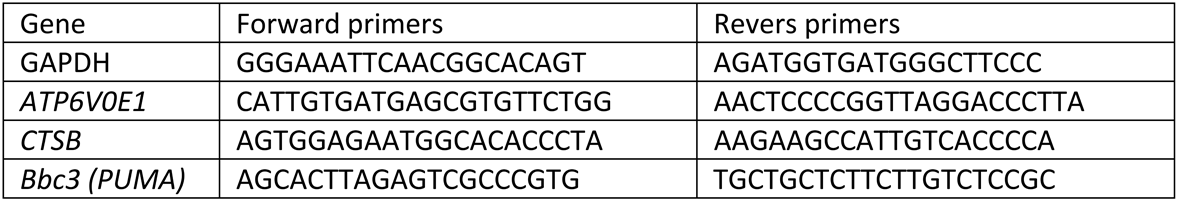

### Incucycte apoptosis assay

Apoptosis was measured by using the Essen BioScience CellPlayer caspase 3/7 reagent according to manufacturer’s instructions. Briefly, 5000 MEFs were seeded in 96-well plates and grown overnight. Caspase 3/7 reagent was added at a 5 µM final concentration to full medium (control cells) or to starvation medium (starved cells). Kinetic activation of caspases was monitored by image acquisition in an IncuCyte FLR equipped with a 20x objective every 4 h. Vybrant DyeCycle Green stain was added directly to the cells after the final scan to determine total cell number. The Incucyte object counting algorithm was used for quantifications. Results are presented as a percentage of apoptotic cells corresponding to ratio between the number of caspase 3/7 positive objects and the total number of DNA containing objects.

### Statistical analyses

Statistical analyses were performed using the Graphpad Prism 6.0 software. Differences between three groups or more were evaluated using multiple t-test or ANOVA test followed by Bonferroni test for multiple comparisons. Comparisons between two groups were performed using the unpaired t-test with Welch’s correction. Data are presented as mean ± SEM. Symbols used are: ns: P > 0.05; *: P ≤ 0.05; **: P ≤ 0.01; *** P ≤ 0.001; **** P ≤ 0.0001.

## ACKNOWLEDGEMENTS

We thank all the members of the Besson and Manenti laboratories for stimulating discussions. We are very grateful to Dr. D.M. Sabatini (Whitehead Institute for Biomedical Research, Cambridge, USA), Dr. J. Debnath (UCSF, San Francisco, USA) and Dr. N. Mizushima (Metropolitan Institute of Medical Science, Tokyo, Japan) for providing reagents. We thank Dr. Vito W. Rebecca (University of Pennsylvania, Philadelphia, USA) for technical advices with PLA experiments. A.N. was supported by studentships from the Ministère de l’Enseignement Supérieur et de la Recherche and from the Fondation ARC pour la Recherche sur le Cancer. J.C. is supported by a studentship from the Region Midi-Pyrénées and Université Paul Sabatier – Toulouse III. R.T.P. is supported by a studentship from the Ligue Nationale Contre le Cancer. S.M. is supported by a grant from the Ligue Nationale Contre le Cancer. This project was supported by funds from the Ligue Nationale Contre le Cancer and Fondation ARC pour la Recherche sur le Cancer to A.B. A.B. is supported by an “FRM Equipes” grant (DEQ20170336707) from the Fondation pour la Recherche Médicale.

