## Supplementary material for "p27 regulates the autophagy-lysosomal pathway via the control of Ragulator and mTOR activity in amino acid deprived cells": Fig S1

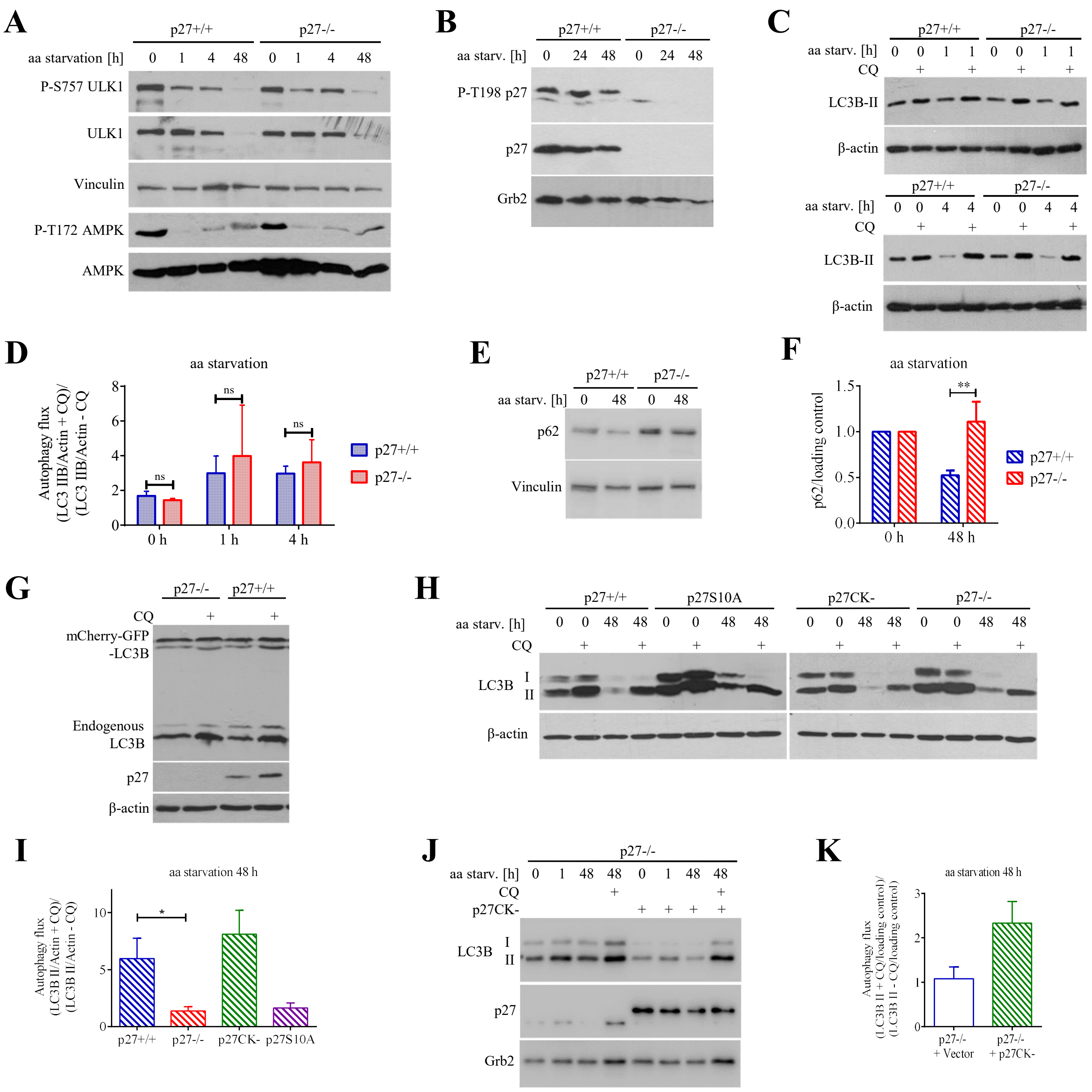


**Figure S1: Cytoplasmic p27 promotes autophagy in a CDK-independent manner in aa deprived cells**

(A) AMPK and ULK1 signaling in starved MEFs. Immunoblots for P-S757 ULK1 and total ULK1 in MEFs aa starved for the indicated times. Vinculin levels were used as loading control. Other membranes of the same extracts were probed for P-T172 AMPK and total AMPK. (B) Phospho-T198 p27 and total p27 immunoblots in p27$^{+/+}$ and p27$^{-/-}$ MEFs starved for the indicated times. Grb2 was used as loading control. (C) LC3B immunoblotting in p27$^{+/+}$ and p27$^{-/-}$ MEFs in full medium (0 h) or aa-deprived for 1 h and 4 h ± 50 µM chloroquine (CQ) for 1 h and 2 h, respectively. β-actin was used as loading control. (D) Quantification of autophagy flux (ratio of LC3B-II/loading control + CQ by LC3B-II/loading control – CQ) from at least 3 independent experiments as in C. (E) Immunoblot for p62 in p27$^{+/+}$ and p27$^{-/-}$ MEFs in full medium (0 h) or aa starved for 48 h. (F) Bar graph shows mean of p62 fold change normalized to 0 h from 7 experiments as shown in E. (G) p27$^{+/+}$ and p27$^{-/-}$ MEFs retrovirally infected with mCherry-eGFP-LC3B ± 50 µM CQ for 2 h. Membranes were probed as indicated. β-actin was used as loading control. (H) LC3B immunoblot in p27$^{+/+}$, p27$^{-/-}$, p27$^{CK-}$ and p27$^{S10A}$ MEFs in full medium (0 h) or aa starved for 48 h ± 50 µM CQ for 2 h. β-actin was used as loading control. p27$^{CK-}$ cells behaved like p27$^{+/+}$ cells, indicating a CDK-independent role of p27, while p27$^{S10A}$ cells in which p27 is sequestered in the nucleus behaved like p27$^{-/-}$ cells, indicating a requirement for cytoplasmic localization of the protein. (I) Quantification of LC3B-II turnover in MEFs aa starved for 48 h as described in H from at least three independent experiments. p27$^{+/+}$ and p27$^{-/-}$ data already appears in Fig 1F. (J) LC3B immunoblot in p27$^{-/-}$ cells infected with either empty vector or p27$^{CK-}$. p27 levels are indicated. Grb2 was used as loading control. (K) Quantification of autophagy flux (ratio of LC3B-II/loading control + CQ by LC3B-II/loading control – CQ) in cells aa deprived for 48 h as in J, normalized to p27$^{-/-}$ + vector condition. Experiment was performed twice. (D, F, I, K) Graphed data are presented as means ± SEM. Statistical significance was evaluated by 2-way ANOVA (D), unpaired t-test with Welch's correction (F) or one-way ANOVA (I). **= $p<0.01$; *=$p<0.05$; ns: $p > 0.05$.
