## Supplementary material for "p27 regulates the autophagy-lysosomal pathway via the control of Ragulator and mTOR activity in amino acid deprived cells": Fig S2

Single antibody control PLA - p27+/+ - aa starvation 18 h

| PLA p27<br>F-Actin<br>Hoechst | PLA LAMP1<br>F-Actin<br>Hoechst | PLA LAMTOR1<br>F-Actin<br>Hoechst | PLA LAMTOR4<br>F-Actin<br>Hoechst |
|---|---|---|---|

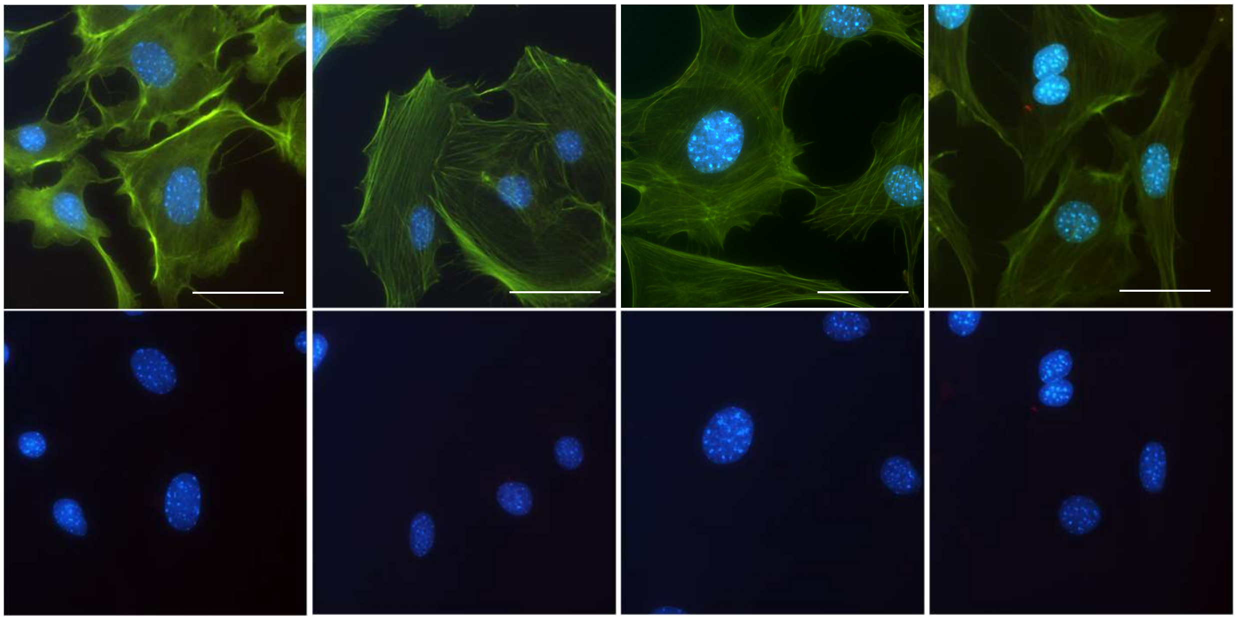

PLA probes only – aa starvation 18 h

p27+/+ p27-/-

PLA
F-Actin
Hoechst

PLA
Hoechst

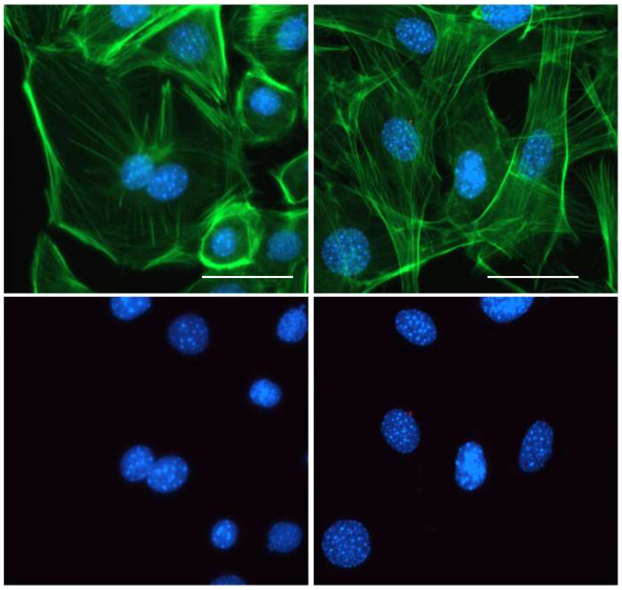

**Figure S2: Proximity ligation assay controls.**
The indicated cells were aa starved for 18 h and submitted to PLA labelling with all the reagents except that only one primary antibody was added (upper panels) or in absence of primary antibodies (lower panels). After the PLA reaction, cells were counterstained for F-actin and nuclear DNA as in the main figures.
