## Supplementary material for "p27 regulates the autophagy-lysosomal pathway via the control of Ragulator and mTOR activity in amino acid deprived cells": Fig S3

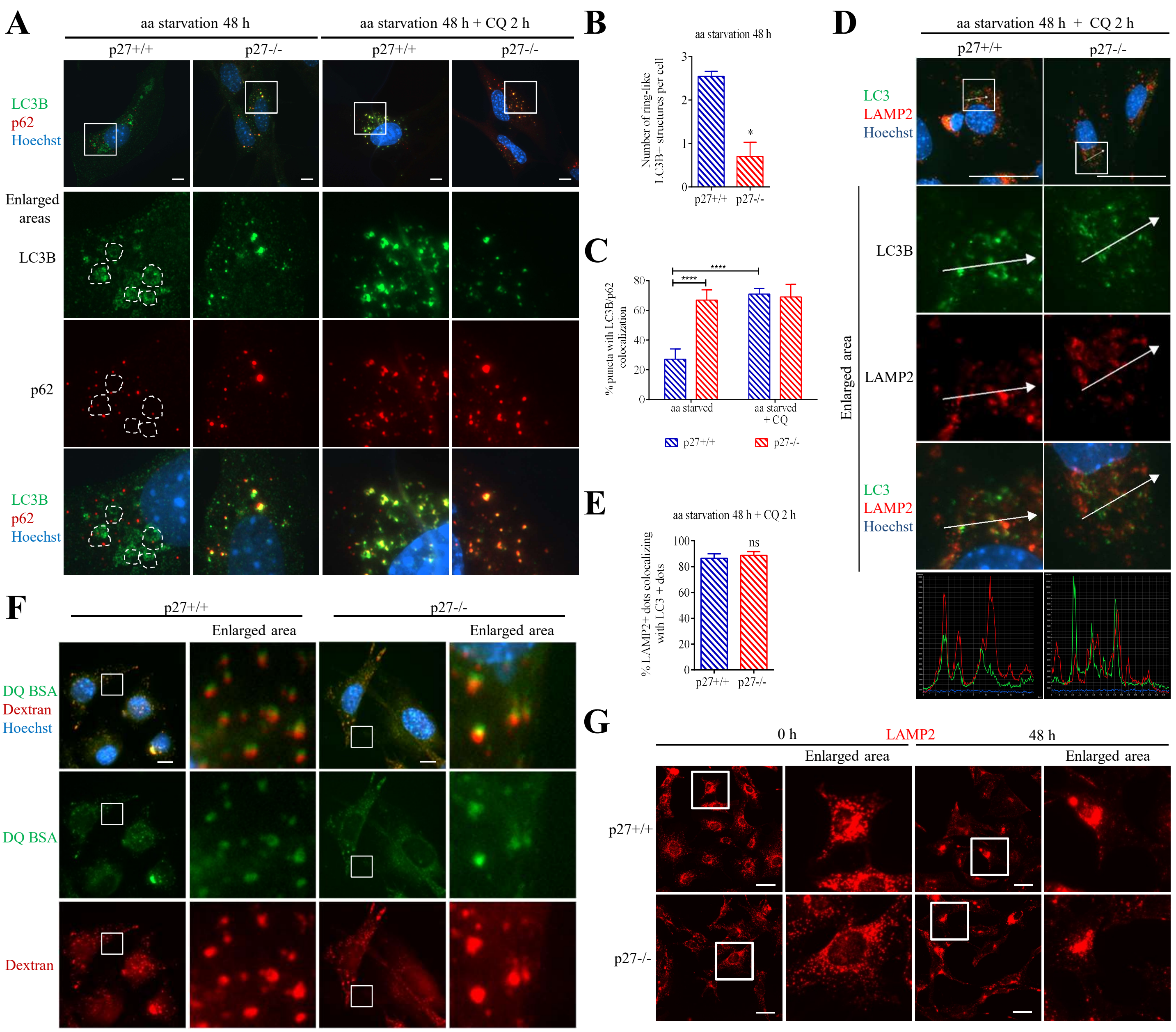


**Figure S3: p27 promotes autophagosome maturation but does not affect autophagogome-lysosome fusion and fluid-phase endocytosis.**

(A) LC3B and p62 immunostaining in p27$^{+/+}$ and p27$^{-/-}$ MEFs aa starved for 48 h ± 50 µM CQ for 2 h. Dotted lines delineate LC3B-positive ring-like structures. (B) Quantification of LC3B-positive ring-like aggregates per cell in MEFs aa deprived for 48 h. At least 10 cells of each genotype were analyzed per experiment (n=3). (C) Quantification of LC3B dots colocalizing with p62 signal in cells from experiments described in A. At least 250 LC3B dots per condition were analyzed in each experiment (n=3). (D) LC3B and LAMP2 immunostaining in p27$^{+/+}$ and p27$^{-/-}$ MEFs aa-starved for 48 h in presence of 50 µM CQ for 2 h. Graphs display the fluorescence intensity (arbitrary unit) in each channel over the distance depicted by the arrows. (E) Quantification of LAMP2 signal overlapping with LC3 dots in cells as described in D. At least 5740 LC3B dots were analyzed per cell line in each experiment (n=3). (F) Representative images of DQ-BSA and Dextran-TRITC in p27$^{+/+}$ and p27$^{-/-}$ MEFs in full medium. Experiment was performed 3 times. (G) LAMP2 immunostaining in cells in full medium (0 h) or aa-starved for 48 h. (H) Quantification of LAMP2 fluorescence intensity in cells from experiments described in G. Values were normalized to p27$^{+/+}$ cells in each condition (n=3). At least 5 images of each genotype, acquired with a 20X objective, were analyzed per condition. (B, C, E, H) Bar graphs shows means ± SEM. Statistical significance was evaluated by unpaired t-test with Welch's corrections (B, E) or by two-way ANOVA test (C, H). ns: $p > 0.05$; *: $p \leq 0.05$; ****: $p \leq 0.0001$. Scale bars are 10 µm.

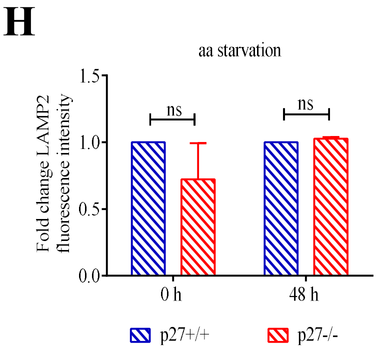
