## Supplementary material for "p27 regulates the autophagy-lysosomal pathway via the control of Ragulator and mTOR activity in amino acid deprived cells": Fig S4

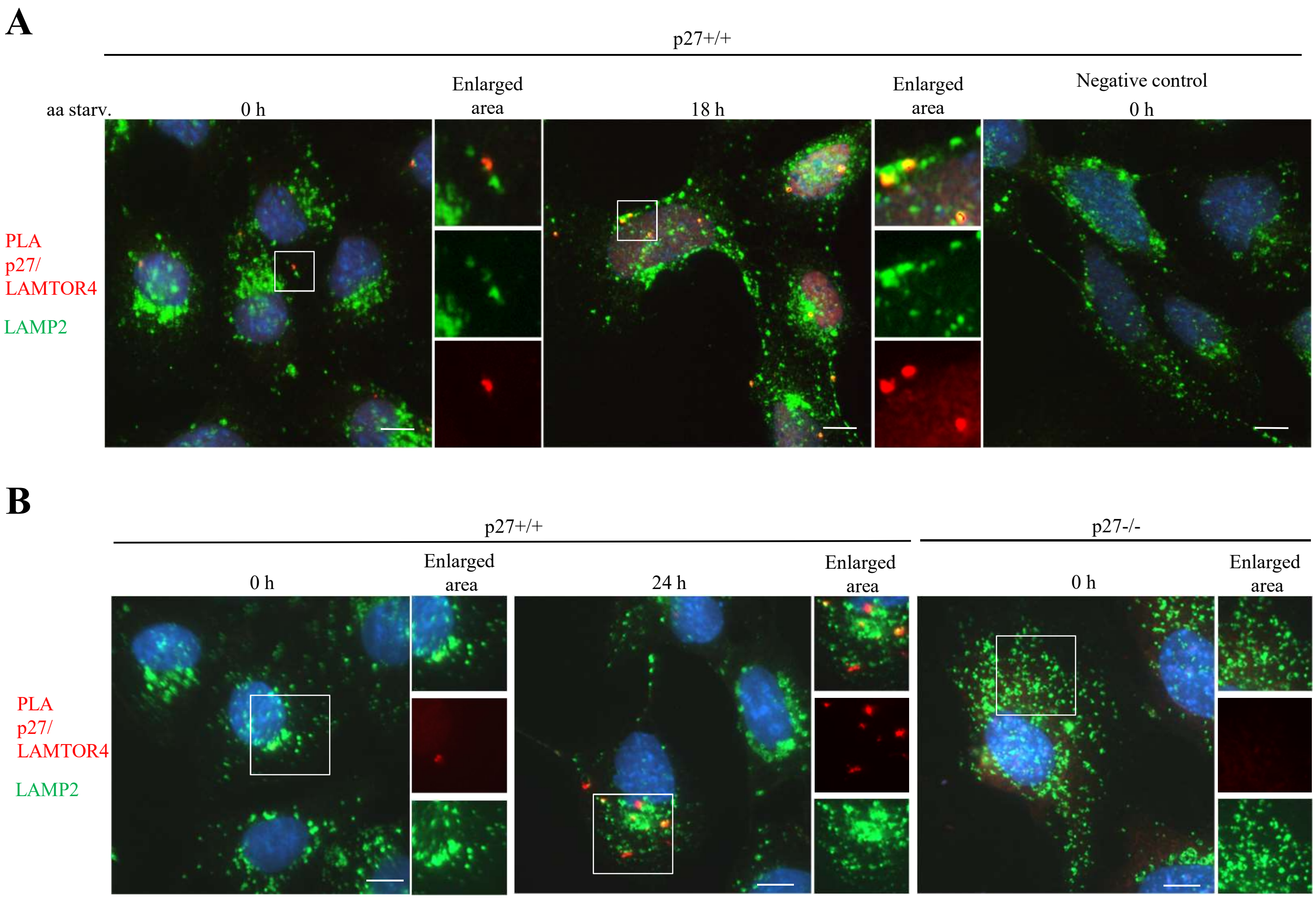


**Figure S4: p27 binds to Ragulator on lysosomes.**
(A) $p27^{+/+}$ MEFs in full medium or aa-deprived for 18 h were subjected to PLA using anti p27 and LAMTOR4 antibodies. Coverslips were then stained for the lysosomal marker LAMP2. PLA probed without primary antibodies were used as negative controls. Representative images from two independent experiments. (B) $p27^{+/+}$ and $p27^{-/-}$ MEFs in full medium or aa starved for 24 h were subjected to PLA and LAMP2 staining as in A. Scale bars are 10 µm. Corresponding single antibody PLA controls and probe only controls are shown in Fig. S3.
