## Supplementary material for "p27 regulates the autophagy-lysosomal pathway via the control of Ragulator and mTOR activity in amino acid deprived cells": Fig S5

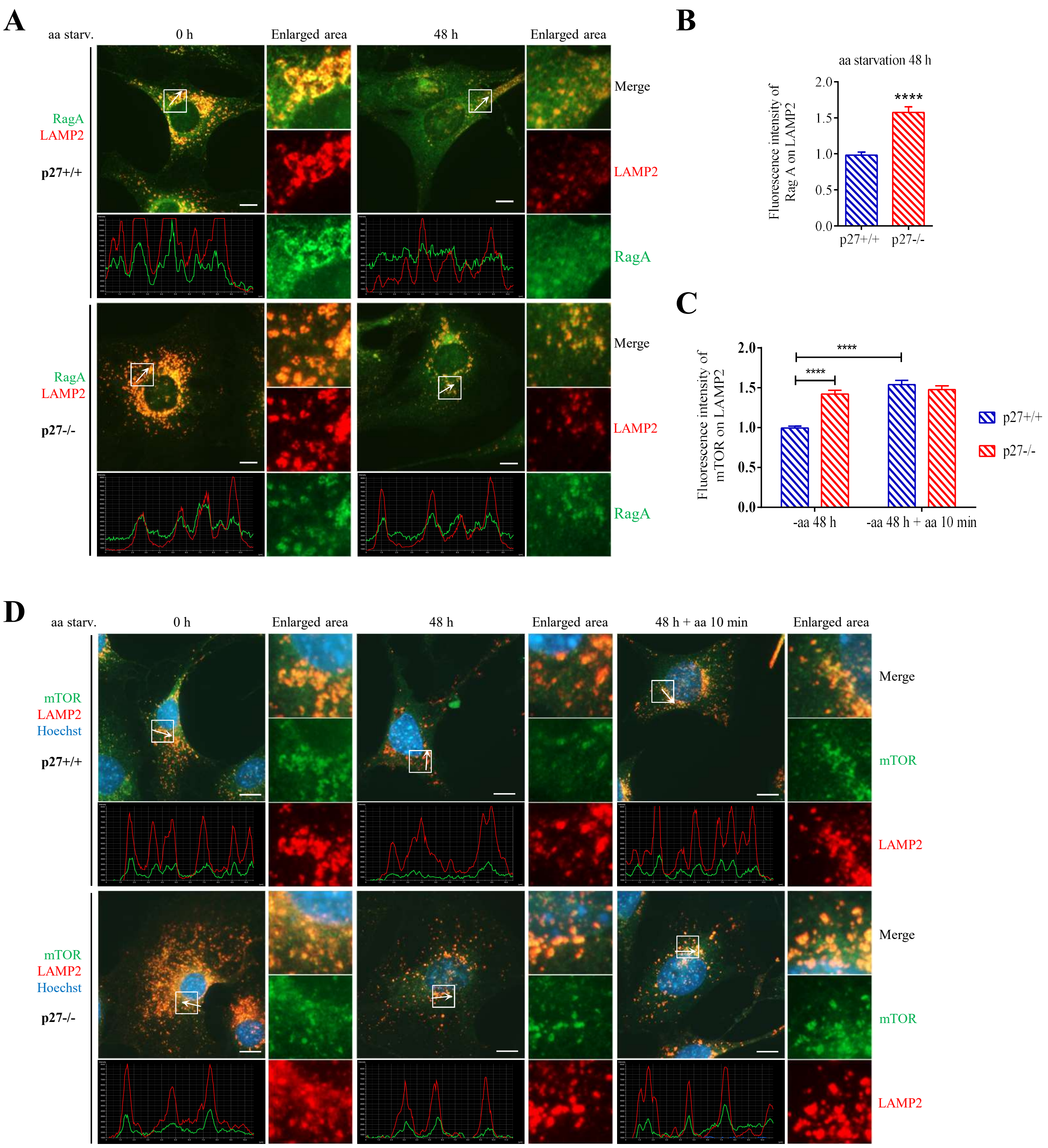


**Figure S5: p27 controls RagA and mTOR recruitment to lysosomes**

(A) RagA and LAMP2 immunostaining in $p27^{+/+}$ and $p27^{-/-}$ MEFs in full medium or aa starved for 48 h. Graphs display the fluorescence intensity (arbitrary unit) in each channel over the distance depicted by the arrows. (B) Quantification of the fluorescence intensity of RagA overlapping with LAMP2 signal normalized to $p27^{+/+}$. Experiment was performed three times. At least 7000 LAMP2+ dots per experiment were analyzed. (C) Quantification of the fluorescence intensity of mTOR overlapping with LAMP2 signal normalized to $p27^{+/+}$ aa-starved for 48 h. At least 3600 LAMP2+ dots were analyzed in each of two experiments. (D) mTOR and LAMP2 immunostaining in $p27^{+/+}$ and $p27^{-/-}$ MEFs either in full medium (0 h), aa starved for 48 h or aa starved for 48 h and re-fed with amino acids for 10 min prior to fixation. Data are presented as means ± SEM. Statistical differences were evaluated by unpaired t-test with Welch's correction (B) or 2-way ANOVA (C). **** $p \leq 0.0001$. In A and D, scale bars are 10 µm.
