## Supplementary material for "p27 regulates the autophagy-lysosomal pathway via the control of Ragulator and mTOR activity in amino acid deprived cells": Fig S6

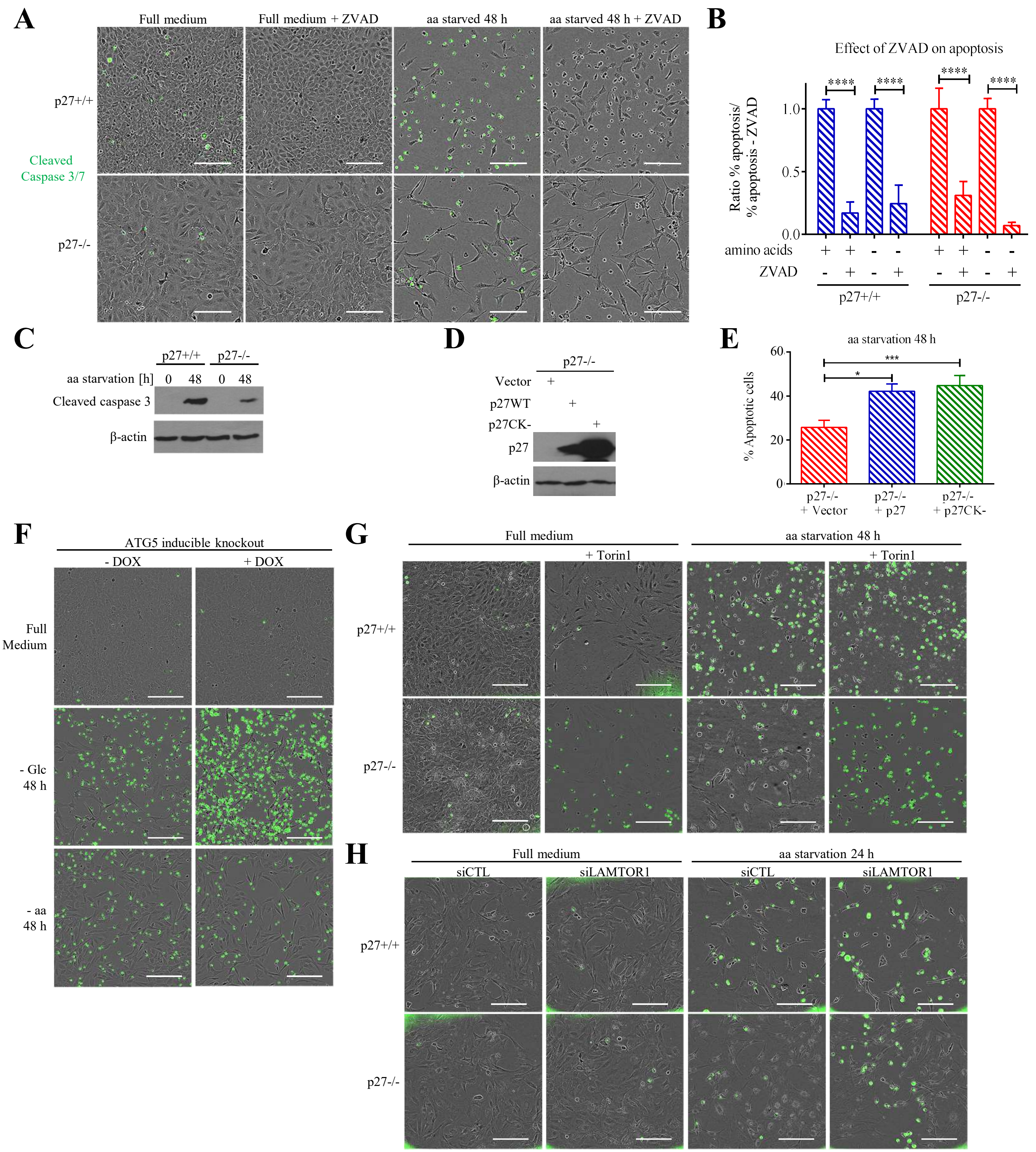


**Figure S6: Elevated mTOR activity in p27$^{-/-}$ cells confers resistance to amino acid starvation-induced apoptosis**

**(A)** Representative Incucyte images of p27$^{+/+}$ and p27$^{-/-}$ MEFs grown in full medium or aa-starved for 48 h ± 20 µM ZVAD caspase inhibitor. Scale bars are 200 µm. **(B)** Quantification of apoptosis in cells from experiments described in B normalized to condition without ZVAD. Bar graph shows means ± SEM from nine experiments. Statistical significance was evaluated by 2-way ANOVA; ****: $p \leq 0.0001$. **(C)** Immunoblot analysis of caspase-3 cleavage in p27$^{+/+}$ and p27$^{-/-}$ MEFs aa-starved for 48 h. β-actin was used as loading control. **(D)** Immunoblot for p27 in p27$^{-/-}$ cells infected with empty vector, p27 or p27$^{CK-}$. β-actin was used as loading control. **(E)** Percentage of apoptosis in cells re-expressing p27 or p27$^{CK-}$ from D aa starved for 48 h. p27 or p27$^{CK-}$ expression restores the susceptibility to apoptosis upon aa starvation. Bar graph shows means ± SEM from at least three experiments. Statistical significance was evaluated by one-way ANOVA; *: $p \leq 0.05$; ***: $p \leq 0.001$. **(F-H)** Representative Incucyte images of phase contrast and green fluorescence representing caspase-3/7 cleavage of (F) ATG5 inducible MEFs ± 10 ng/ml doxycycline in full medium or glucose or aa-starved for 48 h, (G) p27$^{+/+}$ and p27$^{-/-}$ MEFs in full medium or aa-starved for 48 h ± 200 nM Torin1, or (H) p27$^{+/+}$ and p27$^{-/-}$ MEFs transfected with control or LAMTOR1 siRNA in full medium or aa-starved for 24 h. Scale bars are 200 µm.
